# Structural basis of MK-97 positive allosteric modulation at the M_4_ mAChR

**DOI:** 10.64898/2026.05.06.723386

**Authors:** Michaela G. Kaoullas, Jesse I. Mobbs, Ziva Vuckovic, Matthew J. Belousoff, Fu Xiao, Keya Joshi, Jinan Wang, Nicholas Barnes, Vi Pham, Mahmuda Yeasmin, Geoff Thompson, Emma T. van der Westhuizen, Manuela Jörg, Ben Capuano, Andrew B. Tobin, Denise Wootten, Patrick M. Sexton, Radostin Danev, Peter J. Scammells, Yinglong Miao, Arthur Christopoulos, Celine Valant, David M. Thal

## Abstract

Positive allosteric modulators (PAMs) of the M_4_ muscarinic acetylcholine receptor (mAChR) represent a promising therapeutic strategy for treating cognitive deficits and neuropsychiatric disorders. While first-generation M_4_ mAChR PAMs, like LY2033298, demonstrated proof-of-concept, second-generation compounds, such as MK-97, exhibit substantially improved potency and reduced species variability. Here we report the cryo-EM structure of the M_4_ mAChR bound to the endogenous agonist, acetylcholine, and MK-97 at 2.7 Å resolution, revealing the molecular basis for improved M_4_ mAChR PAM activity. MK-97 adopts a distinctive ‘boomerang’-shaped conformation within the extracellular-facing allosteric binding site, with a central pyridine vertex, a lower cyclopentylmethylpyrazole arm extending toward the floor of the orthosteric site, and an upper isoindolinone arm projecting toward extracellular loop 2 (ECL2). This extended binding mode establishes a distributed interaction network across transmembrane helices TM2, TM3, TM5, TM6, and TM7, with key contacts including a hydrogen bond with Y92^2.64^ and a π-π stacking interaction with W435^7.35^. Integration of structural data, molecular dynamics simulations, and mutagenesis validation reveals that the high affinity of MK-97 derives from optimized engagement across all three binding regions rather than dependence on any single critical contact. Insights from comprehensive structure-activity relationship (SAR) studies provide a molecular framework for the rational design of next-generation M_4_ mAChR PAMs with improved pharmacological properties.

**Graphical Abstract:** 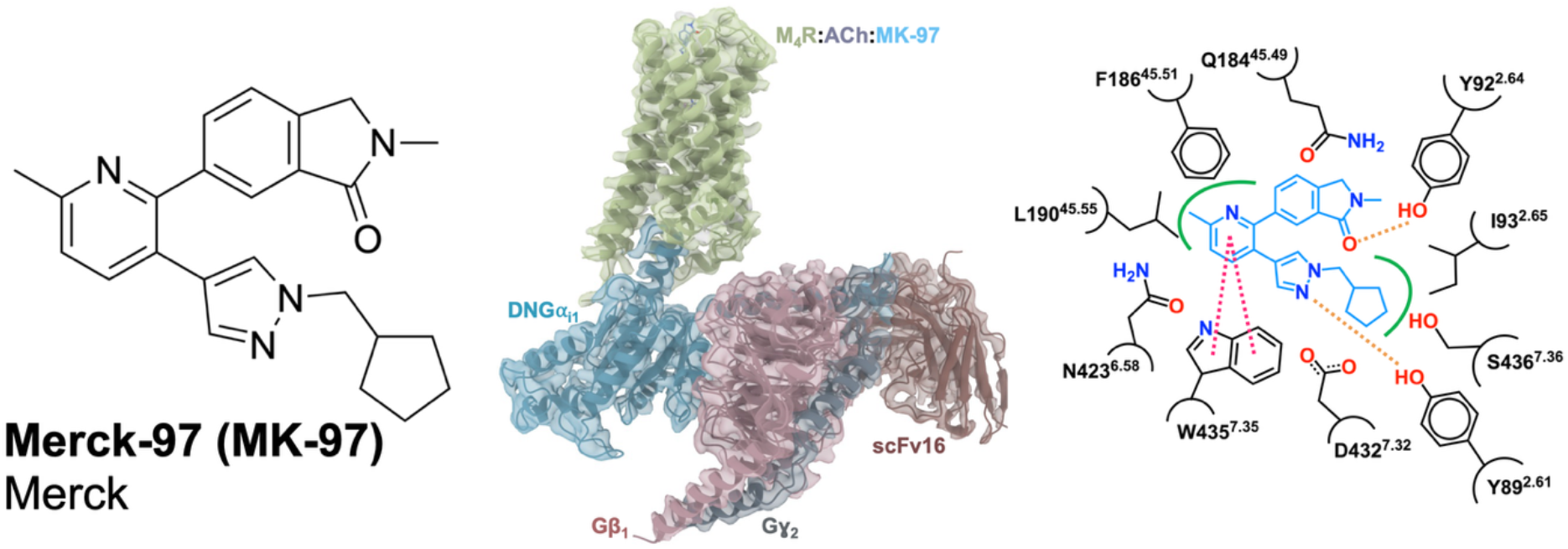

## Introduction

The five muscarinic acetylcholine receptors (M_1_–M_5_ mAChRs) are class A G protein-coupled receptors (GPCRs) that play critical roles in the function of the central and peripheral nervous systems. These receptors mediate the actions of acetylcholine (ACh) and are essential for complex cognitive processes, including learning and memory. Disruption of cholinergic neurotransmission underlies numerous neurological and neuropsychiatric diseases, either directly or through modulation of other vital neurotransmitter systems^1–7^. The M_4_ mAChR subtype has emerged as an attractive therapeutic target due to its high expression in brain regions rich in dopamine receptors, where it modulates dopaminergic signaling involved in cognition and psychosis^7–10^. Extensive preclinical and clinical evidence supports the role of the M_4_ mAChR in CNS disorders, establishing it as a viable therapeutic approach for treating psychosis and cognitive dysfunction in schizophrenia^11–15^. This therapeutic potential was recently validated with the FDA approval of Cobenfy™, a co-formulation of xanomeline (an M_1_/M_4_ mAChR-preferring agonist) and trospium (a peripherally-restricted mAChR antagonist) that demonstrated significant improvements in behavioral dysfunction and cognitive deficits in patients suffering from schizophrenia^16^.

Targeting mAChRs therapeutically has been historically challenging due to the high sequence conservation of the orthosteric ACh binding site across all five subtypes^17–24^. This conservation makes it difficult to develop subtype-selective compounds, as exemplified by xanomeline’s interaction with peripheral M_2_ and M_3_ mAChRs, which causes severe gastrointestinal side effects^25–28^. The co-formulation with trospium was necessary to mitigate these peripheral effects while preserving central therapeutic activity^29,30^. A promising strategy to overcome these limitations involves targeting allosteric sites, which are structurally distinct binding pockets that are less conserved across mAChR subtypes^31,32^. Positive allosteric modulators (PAMs) bind to these sites, enhancing the binding and functional responses of orthosteric ligands via cooperative interactions that stabilize the active conformation of the receptor. This offers the potential for greater mAChR subtype selectivity and an improved clinical safety profile^33,34^. Such selectivity would allow M_4_ mAChR activation without the peripheral side effects that limit orthosteric agonists. Intensive drug discovery efforts have yielded several M_4_ mAChR-selective PAMs (**Figure 1A**), beginning with the first-generation compounds LY2033298 and VU0467154. While these compounds established proof-of-concept for M_4_ mAChR allosteric modulation and demonstrated efficacy in preclinical models of psychosis, their clinical translation was hampered by species variability, poor pharmacokinetic properties, and limited brain penetration^5,11,12,35,36^.

**Figure 1:**
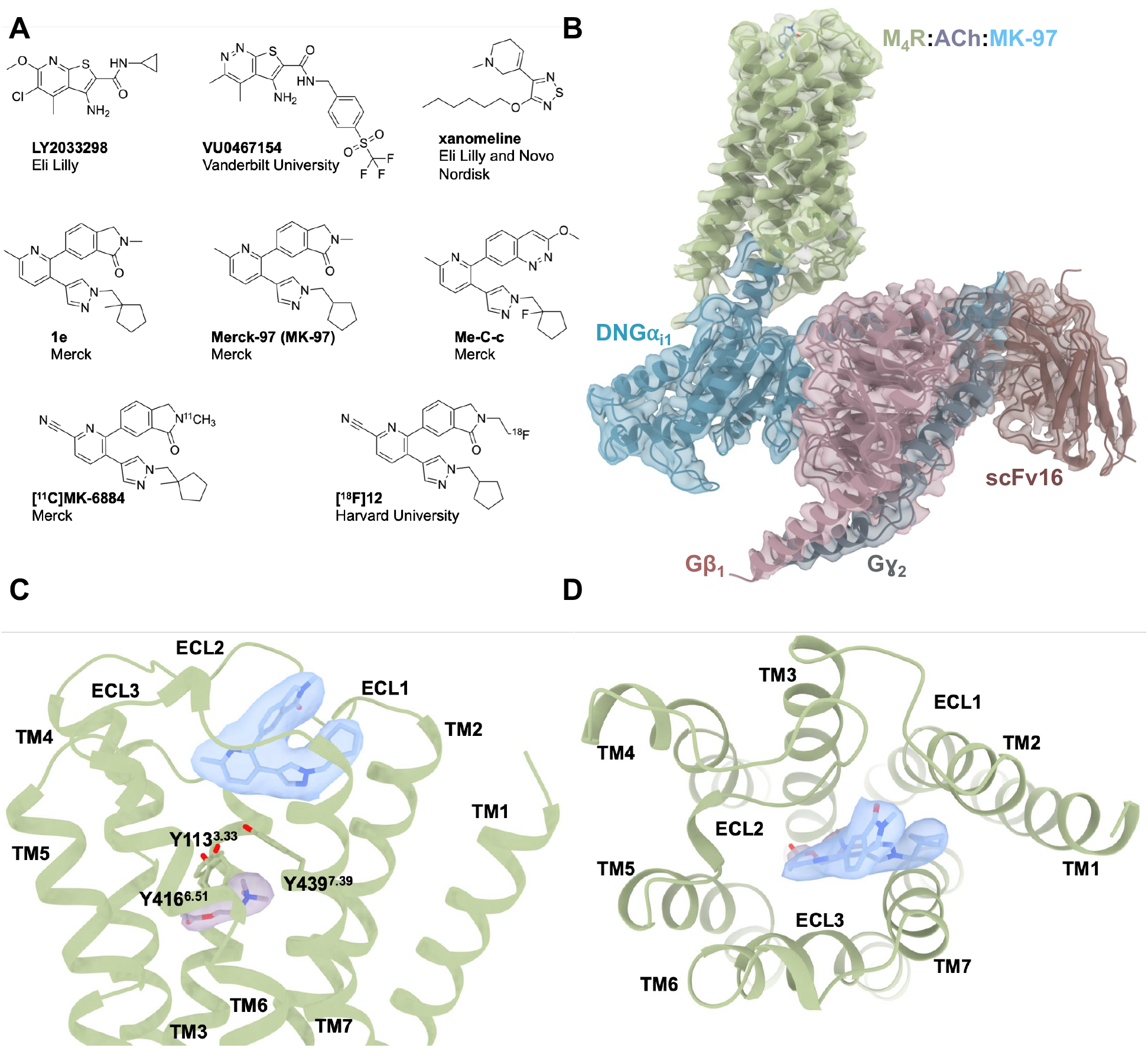
The endogenous agonist, ACh, and PAM, MK-97, bind to the orthosteric and allosteric sites of M_4_ mAChR, respectively. (**A**) Chemical structures of M_4_ mAChR PAMs. (**B**) Membrane view of the consensus cryo-EM map and model of MK-97 bound to M_4_ mAChR in complex with DNGα_i1_, β_1_γ_2_, scFv16. The receptor is shown in green, the heterotrimeric G protein is shown in blue, pink, and grey for the α, β, and γ subunits, respectively, and scFv16 in brown. (**C**) Cryo-EM map (contour level 0.324) of ACh, shown in purple, in the orthosteric binding site, and of MK-97, shown in blue, within the allosteric site. The two distinct sites of the M_4_ mAChR are separated by the tyrosine lid, shown as green sticks. (**D**) Extracellular view of the ECV occupied by MK-97. Our cryo-EM maps revealed unambiguous density for MK-97 in the ECV (**Figure 1C**,**D**), the ‘common’ allosteric site across the mAChR family^32^. The ECV allosteric site is defined by a floor comprising the tyrosine lid and residues from TM2, TM6, TM7, ECL2, and ECL3. These regions undergo significant movements during receptor activation, resulting in ECV contraction that is further stabilized by allosteric ligand binding^17,22^. MK-97 adopts a ‘boomerang’-shaped conformation in the ECV with distinct pendant groups: the isoindolinone (upper arm), pyridine (central vertex), and cyclopentylmethylpyrazole (lower arm) (**Figure 2A**).

The development of second-generation PAMs based on 2,3,6-trisubstituted pyridine scaffolds marked a significant advancement^37^. These compounds, exemplified by molecules containing a pyrazol-4-ylpyridine core structure, display sub-micromolar affinity for the M_4_ mAChR allosteric site alongside high potency, improved receptor subtype selectivity, and better brain penetration. Notable examples include Me-C-c^37,38^ and the PET tracers, [^11^C]MK-6884 and [^18^F]12^39–43^. Importantly, these second-generation PAMs exhibit approximately 10-fold greater affinity compared to first-generation compounds^42–44^. Understanding the molecular basis for this dramatic improvement in affinity can be aided by high-resolution structural information. Recent advances in cryo-electron microscopy (cryo-EM) have provided structural insights into allosteric modulation of the M_4_ mAChR. Previous structural studies with first-generation PAMs revealed the molecular basis for PAM binding and established the structural foundation for species selectivity, a key challenge in clinical translation^22^. These studies identified key residues in the extracellular vestibule (ECV) that forms the best-characterized mAChR allosteric binding site and undergo conformational changes upon PAM binding. However, the structural basis for the improved affinity of second-generation PAMs remains unknown.

With the emergence of structurally and pharmacologically distinct pyrazol-4-ylpyridine-based PAMs, there is an opportunity to apply structural biology techniques to understand the molecular determinants of their affinity and pharmacological properties. Here, we determined a 2.7 Å cryo-EM structure of the M_4_ mAChR in complex with its cognate Gα_i1_ heterotrimer, co-bound to ACh and the second-generation PAM, Merck-97 (MK-97), a high-affinity pyrazol-4-ylpyridine derivative. This structural information guided molecular dynamics (MD) simulations to probe ligand-receptor interactions over time and targeted mutagenesis studies to investigate the molecular mechanisms underlying the binding and functional properties of MK-97. Our findings reveal a binding mode for MK-97 that differs significantly from first-generation PAMs, providing new insights into M_4_ mAChR allosteric modulation. These findings explain prior structure-activity relationships (SAR) of pyrazol-4-ylpyridine-based PAMs and provide new opportunities for designing next-generation M_4_ mAChR PAMs with enhanced selectivity and clinical potential^42,43^.

## Results

### Structure determination and analysis of the ACh and MK-97-bound M_4_ mAChR

To determine the structure of the M_4_ mAChR co-bound to ACh and MK-97, we applied methodology similar to previously reported M_4_ mAChR complex structures (**Supplementary Figure 1)**^20,22^. The resulting structure was determined to a global resolution of 2.7 Å, providing sufficient cryo-EM maps to model all components of the M_4_ mAChR-DNGα_i1_β_1_γ_2_ complex and the bound ligands (**Figure 1B**). The M_4_R-ACh-MK-97 complex adopts an active conformation with root mean squared deviation (RMSD) values of 0.42–0.48 relative to other agonist-bound M_4_ mAChR structures^22,45^ and displays the canonical features of class A GPCR activation, including TM6 outward movement and rearrangement of conserved activation motifs (**Supplementary Figure 2, 3A**,**B)**^46–50^. As expected, ACh occupies the orthosteric site within the transmembrane bundle, stabilized by conserved aromatic residues and the tyrosine lid (Y113^3.33^, Y416^6.51^, and Y439^7.39^) that contracts during activation (**Figure 1C**)^22,51^.

**Figure 2:**
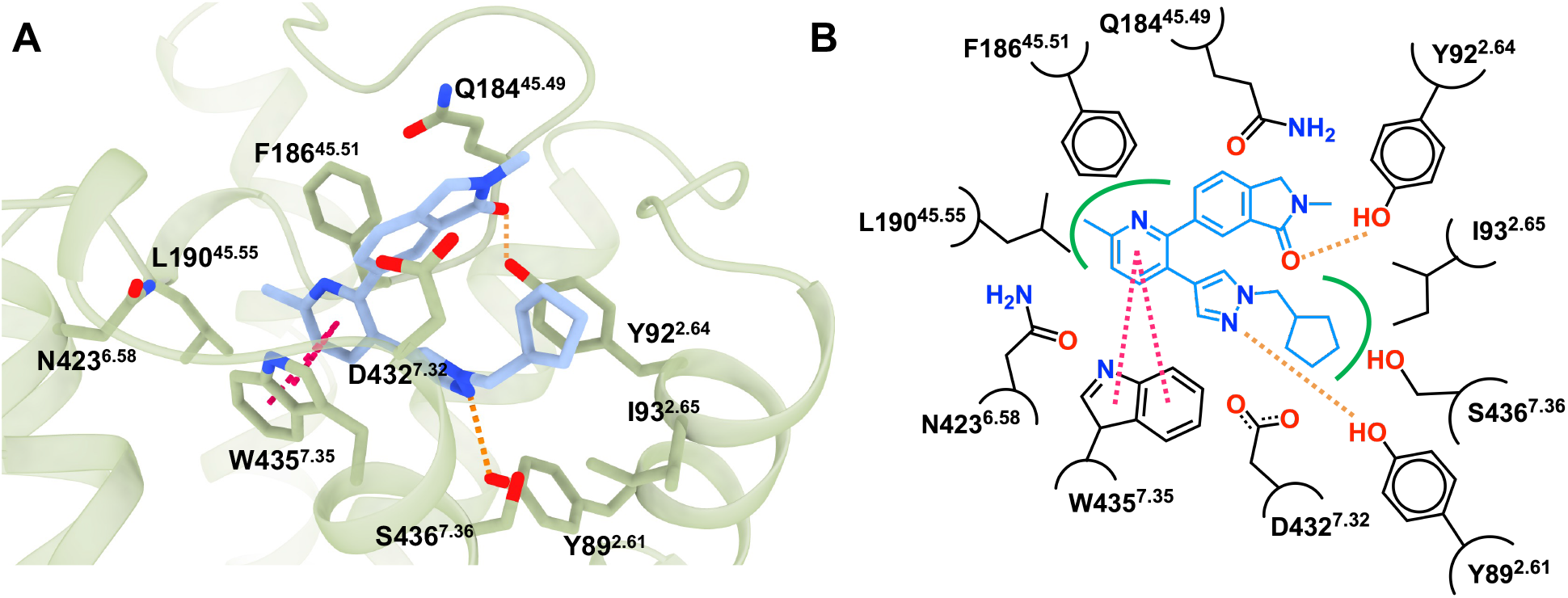
Analysis of the allosteric binding site. (**A**) Interactions between allosteric site residues and MK-97, shown in blue sticks. Polar interactions are indicated by the orange dashed line and π-interactions are shown by magenta dashed lines. (**B**) 2D chemical interactions of MK-97 with M_4_ mAChR residues. Color coding is the same as (**A**) with hydrophobic interactions indicated by the green line.

**Figure 3:**
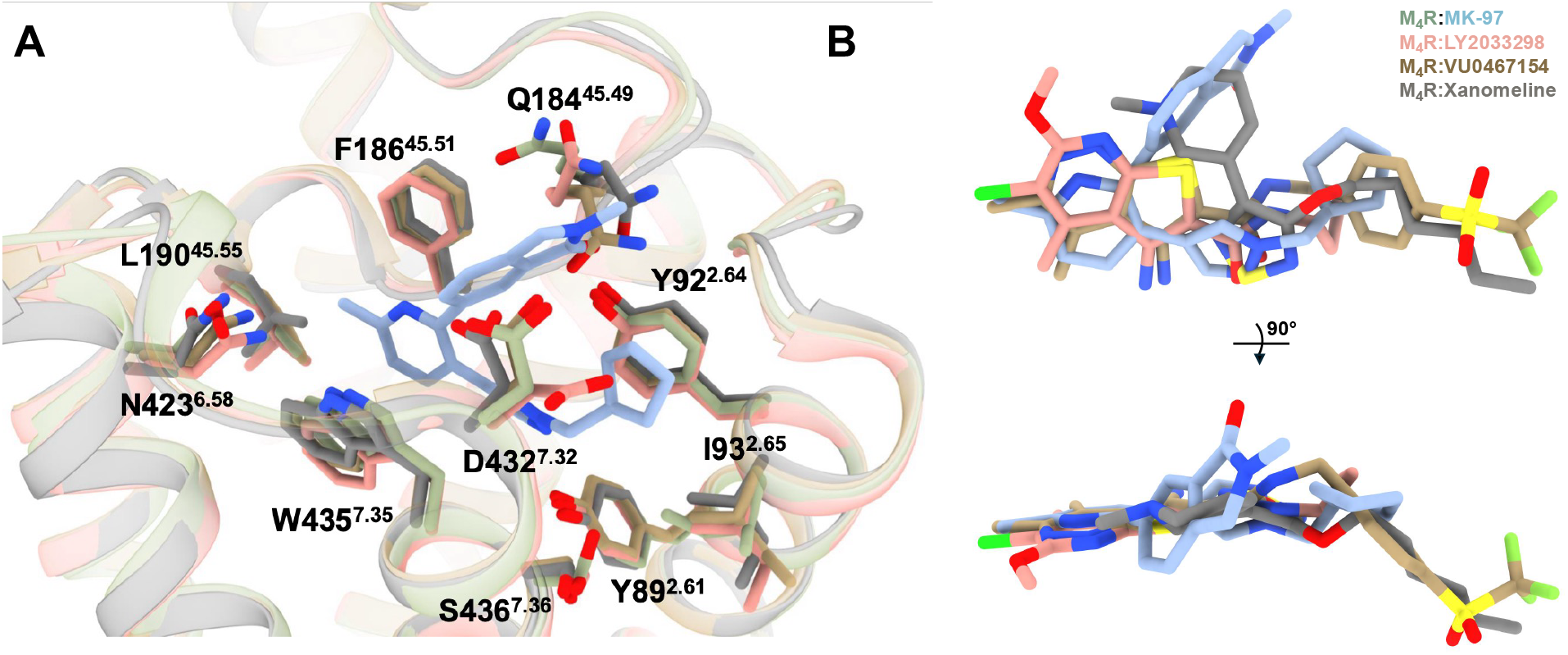
Comparison of the M_4_ mAChR allosteric site. (**A**) Alignment of M_4_ mAChR structures bound to MK-97, LY2033298 (PDB: 7TRP), VU0567154 (PDB: 7TRQ), and xanomeline (PDB: 8FX5). Allosteric site residues of the MK-97 bound M_4_ mAChR are shown as green sticks, residues of the LY2033298, VU0467154, and xanomeline bound M_4_ mAChRs are shown as pink, brown, and grey sticks, respectively. (**B**) Overlay of M_4_ allosteric ligands MK-97, LY2033298, VU0467154, and xanomeline.

The central pyridine vertex forms conserved interactions with the allosteric binding pocket. The pyridine of MK-97 forms a π-π interaction with W435^7.35^, an interaction common to all M_4_ mAChR PAMs and previously shown to be essential for PAM binding (**Figure 2A,B**)^22^. Surrounding the pyridine are hydrophobic residues L190^45.55^, F186^45.51^, Y416^6.51^, and N423^6.58^, which move toward the TM core during activation to form a tight hydrophobic pocket around the central scaffold. This region overlaps with the thieno[2,3-*b*]pyridine core of first-generation PAMs LY2033298 and VU0467154 (**Figure 2A,B**).

The lower cyclopentylmethylpyrazole arm extends into the deeper ECV pocket. The pyrazole nitrogen forms a hydrogen bond with Y89^2.61^, which rotates outward from its position in other agonist-bound structures to accommodate this group **(Figure 2A,B)**. This interaction has been pharmacologically validated in previous mutagenesis studies, with Y89^2.61^A mutations abolishing PAM activity^22,52^. The cyclopentylmethyl group is stabilized by hydrophobic contacts with I93^2.65^, D432^7.32^, and S436^7.36^, with the latter two residues undergoing appreciable movement between inactive and active states. This lower region partially overlaps with the binding poses of LY2033298 and VU0467154 **(Figure 2A,B)**.

The isoindolinone pendant adopts a unique orientation extending toward extracellular loop 2 (ECL2). Unlike first-generation PAMs, the isoindolinone extends outward toward ECL2, causing slight displacement and rotation of Q184^45.49^ (ECL residues are numbered 45.X, denoting their position between TM4 and TM5, with 45.50 being the conserved cysteine)^53^ to position the isoindolinone between ECL1 and ECL2 **(Figure 3A,B)**. The carbonyl oxygen of the isoindolinone forms a hydrogen bond with Y92^2.64^. When comparing the binding pose of MK-97 to that of xanomeline in the M_4_ mAChR allosteric site, the upper the upper arm of MK-97 is positioned over the thiadiazole of xanomeline and overlaps with the tetrahydropyridine ring of xanomeline **(Figure 3A,B)**.

Comparison to first-generation PAMs reveals a distinct allosteric binding mode for MK-97. While MK-97 shares conserved interactions with first-generation PAMs, particularly the W435^7.35^ π-π stacking interaction and the Y89^2.61^ hydrogen bond, notable differences are also evident. Most strikingly, F186^45.51^ does not form π-stacking interactions with MK-97, in contrast to its engagement with LY2033298, VU0467154, and xanomeline in previously-solved structures^17,20,22,45^. The F186A^45.51^ mutation abolishes affinity for these first-generation PAMs^22^, yet has minimal impact on MK-97, whose binding extends into a distinct pocket created by the isoindolinone interaction with ECL2. This unique binding mode, particularly the upper arm extension into the ECL2 region, provides a structural rationale for the enhanced affinity of second-generation PAMs, and thus guided our subsequent mutagenesis and functional studies.

### Validation of the allosteric binding mode of MK-97

Prior mutagenesis studies identified residues at the orthosteric and allosteric sites that contribute to key pharmacological parameters including PAM affinity (p*K*_B_), binding cooperativity with the orthosteric ligand (*α*), direct allosteric ligand signaling efficacy (*τ*_B_), and allosteric modulation of orthosteric ligand efficacy (*β*)^17,52,54^. These studies revealed a network of conserved and non-conserved M_4_ mAChR residues spanning both sites that mediate allosteric signaling transmission. Given the novel binding pose of MK-97, we performed equilibrium radioligand binding experiments at wild-type (WT) and mutant M_4_ mAChRs to validate structural observations and delineate the molecular determinants of MK-97 affinity and cooperativity with ACh. We used interaction competition binding assays with the orthosteric antagonist [^3^H]-NMS and ACh in the absence and presence of increasing concentrations of MK-97. An allostery ternary complex model was fitted to the data to determine MK-97 ligand affinity (p*K*_B_) for the free receptor (i.e., affinity for the allosteric site in the absence of orthosteric ligand) and binding cooperativity (*α*) factors quantifying the magnitude of change in affinity in the presence of co-bound orthosteric ligand (**Figure 4, Table 1**).

**Table 1:**
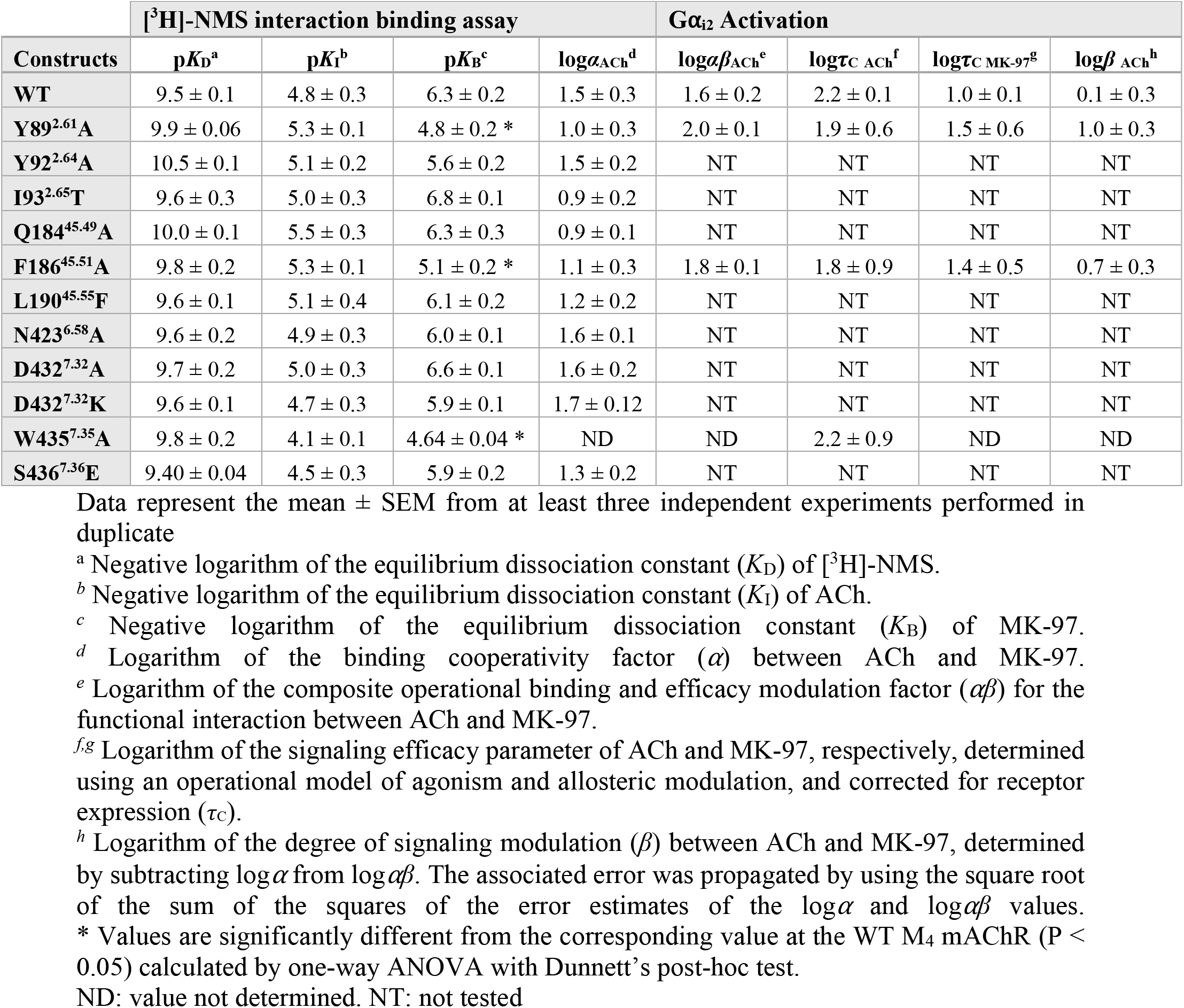
Pharmacological parameters from radioligand binding and functional experiments.

| Constructs | [<sup>3</sup>H]-NMS interaction binding assay | | | | G$\alpha_{i2}$ Activation | | | |
| --- | --- | --- | --- | --- | --- | --- | --- | --- |
| | pK<sub>D</sub><sup>a</sup> | pK<sub>I</sub><sup>b</sup> | pK<sub>B</sub><sup>c</sup> | logα<sub>ACh</sub><sup>d</sup> | log$\alpha\beta$<sub>ACh</sub><sup>e</sup> | log$\tau_C$<sub>ACh</sub><sup>f</sup> | log$\tau_C$<sub>MK-97</sub><sup>g</sup> | logβ<sub>ACh</sub><sup>h</sup> |
| WT | 9.5 ± 0.1 | 4.8 ± 0.3 | 6.3 ± 0.2 | 1.5 ± 0.3 | 1.6 ± 0.2 | 2.2 ± 0.1 | 1.0 ± 0.1 | 0.1 ± 0.3 |
| Y89<sup>2,61</sup>A | 9.9 ± 0.06 | 5.3 ± 0.1 | 4.8 ± 0.2 * | 1.0 ± 0.3 | 2.0 ± 0.1 | 1.9 ± 0.6 | 1.5 ± 0.6 | 1.0 ± 0.3 |
| Y92<sup>2,64</sup>A | 10.5 ± 0.1 | 5.1 ± 0.2 | 5.6 ± 0.2 | 1.5 ± 0.2 | NT | NT | NT | NT |
| I93<sup>2,65</sup>T | 9.6 ± 0.3 | 5.0 ± 0.3 | 6.8 ± 0.1 | 0.9 ± 0.2 | NT | NT | NT | NT |
| Q184<sup>45,49</sup>A | 10.0 ± 0.1 | 5.5 ± 0.3 | 6.3 ± 0.3 | 0.9 ± 0.1 | NT | NT | NT | NT |
| F186<sup>45,51</sup>A | 9.8 ± 0.2 | 5.3 ± 0.1 | 5.1 ± 0.2 * | 1.1 ± 0.3 | 1.8 ± 0.1 | 1.8 ± 0.9 | 1.4 ± 0.5 | 0.7 ± 0.3 |
| L190<sup>45,55</sup>F | 9.6 ± 0.1 | 5.1 ± 0.4 | 6.1 ± 0.2 | 1.2 ± 0.2 | NT | NT | NT | NT |
| N423<sup>6,58</sup>A | 9.6 ± 0.2 | 4.9 ± 0.3 | 6.0 ± 0.1 | 1.6 ± 0.1 | NT | NT | NT | NT |
| D432<sup>7,32</sup>A | 9.7 ± 0.2 | 5.0 ± 0.3 | 6.6 ± 0.1 | 1.6 ± 0.2 | NT | NT | NT | NT |
| D432<sup>7,32</sup>K | 9.6 ± 0.1 | 4.7 ± 0.3 | 5.9 ± 0.1 | 1.7 ± 0.12 | NT | NT | NT | NT |
| W435<sup>7,35</sup>A | 9.8 ± 0.2 | 4.1 ± 0.1 | 4.64 ± 0.04 * | ND | ND | 2.2 ± 0.9 | ND | ND |
| S436<sup>7,36</sup>E | 9.40 ± 0.04 | 4.5 ± 0.3 | 5.9 ± 0.2 | 1.3 ± 0.2 | NT | NT | NT | NT |

Data represent the mean ± SEM from at least three independent experiments performed in duplicate
<sup>a</sup> Negative logarithm of the equilibrium dissociation constant ( $K_D$ ) of [<sup>3</sup>H]-NMS.
<sup>b</sup> Negative logarithm of the equilibrium dissociation constant ( $K_I$ ) of ACh.
<sup>c</sup> Negative logarithm of the equilibrium dissociation constant ( $K_B$ ) of MK-97.
<sup>d</sup> Logarithm of the binding cooperativity factor ( $\alpha$ ) between ACh and MK-97.
<sup>e</sup> Logarithm of the composite operational binding and efficacy modulation factor ( $\alpha\beta$ ) for the functional interaction between ACh and MK-97.
<sup>f,g</sup> Logarithm of the signaling efficacy parameter of ACh and MK-97, respectively, determined using an operational model of agonism and allosteric modulation, and corrected for receptor expression ( $\tau_C$ ).
<sup>h</sup> Logarithm of the degree of signaling modulation ( $\beta$ ) between ACh and MK-97, determined by subtracting $\log \alpha$ from $\log \alpha\beta$ . The associated error was propagated by using the square root of the sum of the squares of the error estimates of the $\log \alpha$ and $\log \alpha\beta$ values.
\* Values are significantly different from the corresponding value at the WT M<sub>4</sub> mAChR (P < 0.05) calculated by one-way ANOVA with Dunnett's post-hoc test.

**Figure 4:**
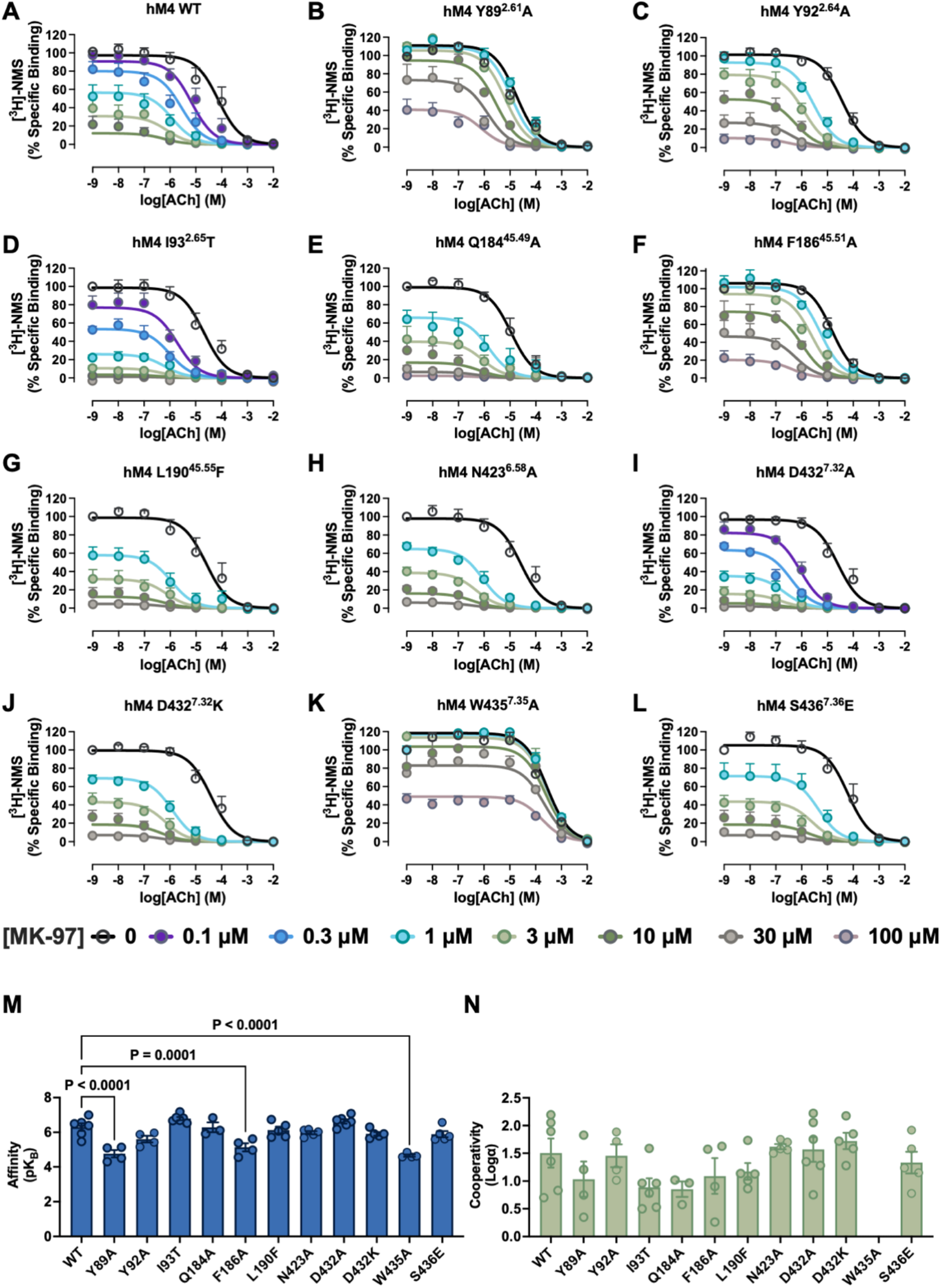
Pharmacological characterization of MK-97 in radioligand binding at WT and mutant M_4_ mAChRs. (**A-L**) Intact CHO cells [^3^H]-NMS inhibition binding curves with ACh in the presence or absence of MK-97 at (A) WT and (B-L) mutant M_4_ mAChRs. All data are the mean ± SEM from at least three independent experiments performed in duplicate. Curve fits represent the global fit of the data to an allosteric ternary complex model. For W435A logα was constrained to zero. **(M)** Negative logarithm of the binding affinity of MK-97 (p*K*_B_) at receptor in the absence of orthosteric ligand. **(N)** Logarithm of the binding cooperativity (logα) between ACh and MK-97. *Values are significantly different from human WT M_4_ mAChR (P < 0.05) calculated by one-way ANOVA with Dunnett’s post-hoc test. ND: value not determined.

At the WT M_4_ mAChR, MK-97 had a p*K*_B_ of 6.3 ± 0.2 (i.e., K_B_ ~500 nM), consistent with previous reports^43,55^, and positively modulated ACh binding with a cooperativity factor of log *α* = 1.5 ± 0.2 (i.e., *α* ~ 30; **Table 1**). As commonly observed for interactions between PAMs and antagonists/inverse agonists, MK-97 and [^3^H]-NMS exhibited very high negative cooperativity (*α* → 0), as PAMs stabilize active-state conformations that reduce antagonist affinity at the orthosteric site. Mutation of W435A^7.35^, which forms the central π-π stacking interaction with the pyridine core of MK-97, caused a ~100-fold decrease in PAM affinity and a complete loss of cooperativity with ACh (**Table 1**). The interaction with the tryptophan residue at position 7.35 is shared by all M_4_ mAChR PAMs that have been characterized to date in mutagenesis studies^22^, serving as an ‘anchor’ for the ligand and facilitating conformational changes in ECL3 that stabilize the active state.

Given the extended binding pose of MK-97 toward ECL2, we also assessed the effect of alanine substitutions of Q184^45.49^, F186^45.51^, and L190^45.55^ (**Table 1**). The F186A^45.51^ mutation significantly reduced, but did not abolish, MK-97 affinity (**Figure 4F,M**), contrasting sharply with complete loss of binding affinity for LY2033298, VU0467154, and xanomeline^28,48^. This difference accords with our structural observation that MK-97 lacks π-stacking interactions with F186^45.51^, which are a common feature of first-generation PAM structures, and suggests that the isoindolinone’s extension toward ECL2 provides compensatory stabilizing contacts. The L190F^45.55^ mutation, which introduces the equivalent residue at the M_2_ mAChR (i.e., F181^45.55^), had no effect on the affinity of MK-97 (**Figure 4G**), consistent with MK-97’s demonstrated activity at the M_2_ mAChR^43,45^. Residue Q184^45.49^ appears to interact with the upper isoindolinone arm of MK-97, however, mutation to alanine produced only a minor loss in binding cooperativity with ACh, suggesting that the residue primarily functions in positioning ECL2 rather than directly stabilizing MK-97 binding.

The Y89A^2.61^ mutation caused significant PAM affinity loss, confirming the importance of the hydrogen bond between the MK-97 pyrazole nitrogen and Y89^2.61^ observed in our structure (**Figure 4B,M, Table 1**). Notably, binding cooperativity with ACh remained intact at this mutant, contrasting with its effects on first-generation PAMs^22,52^ and suggesting that the extended receptor residues interaction network of MK-97 compensates for the absence of this side chain. The I93T^2.65^ mutation (corresponding to M_2_ mAChR T84^2.65^) produced only a slight, non-significant increase in binding affinity (**Figure 4D,M**), similar to the slight increase in binding cooperativity observed for LY2033298 at this position^52^. Interestingly, mutation of Y92A^2.64^, which forms a hydrogen bond with the isoindolinone carbonyl in our structure, had no effect on PAM affinity or cooperativity with ACh, suggesting that this interaction is not significant or that other contacts compensate for its loss (**Figure 4C**). Mutation of N423A^6.58^, implicated in cooperativity transmission for first-generation PAMs^52^, did not affect MK-97 affinity or its ability to allosterically modulate ACh (**Figure 4H**). Similarly, mutations of non-conserved TM7 residues D432K^7.32^ and S436E^7.36^ (to their M_3_ and M_1_ mAChR equivalents, respectively) had no impact on MK-97 pharmacology (**Figure 4J,L**), despite these residues undergoing conformational changes during activation^17,22^.

Collectively, the mutagenesis data validate the MK-97 binding mode observed in our structure and reveal key differences from first-generation PAMs. The extended binding pose of MK-97 toward ECL2 reduces dependence on F186^45.51^ π-stacking while maintaining the conserved W435^7.35^ interaction, and introduces new contacts via the isoindolinone arm. This distinct interaction profile, combined with retention of positive cooperativity with ACh at several single-point mutants, suggests that second-generation PAMs may achieve overall enhanced affinity through a more distributed interaction network.

### Functional characterization reveals distinct roles for key binding residues

The binding studies identified W435^7.35^, Y89^2.61^, and F186^45.51^ as critical determinants of MK-97 binding and cooperativity. To assess whether these residues also influence receptor signaling, we used the BRET TruPath^56^ assays measure Gα_i1_ activation at WT and mutant M_4_ mAChRs (**Figure 5A-D**). By applying an operational model of allosterism and agonism^34,54,57^, we quantified the signaling efficacy of ACh (*τ*_A_) and MK-97 (*τ*_B_), as well as the overall magnitude of functional allosteric modulation (*αβ*) imparted by MK-97 (**Table 1**). At the WT M_4_ mAChR, MK-97 exhibited robust positive allosteric modulation of ACh signaling efficacy and demonstrated significant allosteric agonism. The efficacy modulation parameter (*β*) was determined by comparing functional modulation (*αβ*) to binding cooperativity (*α*), yielding a log *β* = 0.09 ± 0.30 at WT M_4_ mAChR (i.e., *β* ~1.2). This near neutral *β* value indicates that the primary allosteric effect of MK-97 is mediated through enhancement of ACh binding affinity, rather than modulation of ACh signaling efficacy.

**Figure 5:**
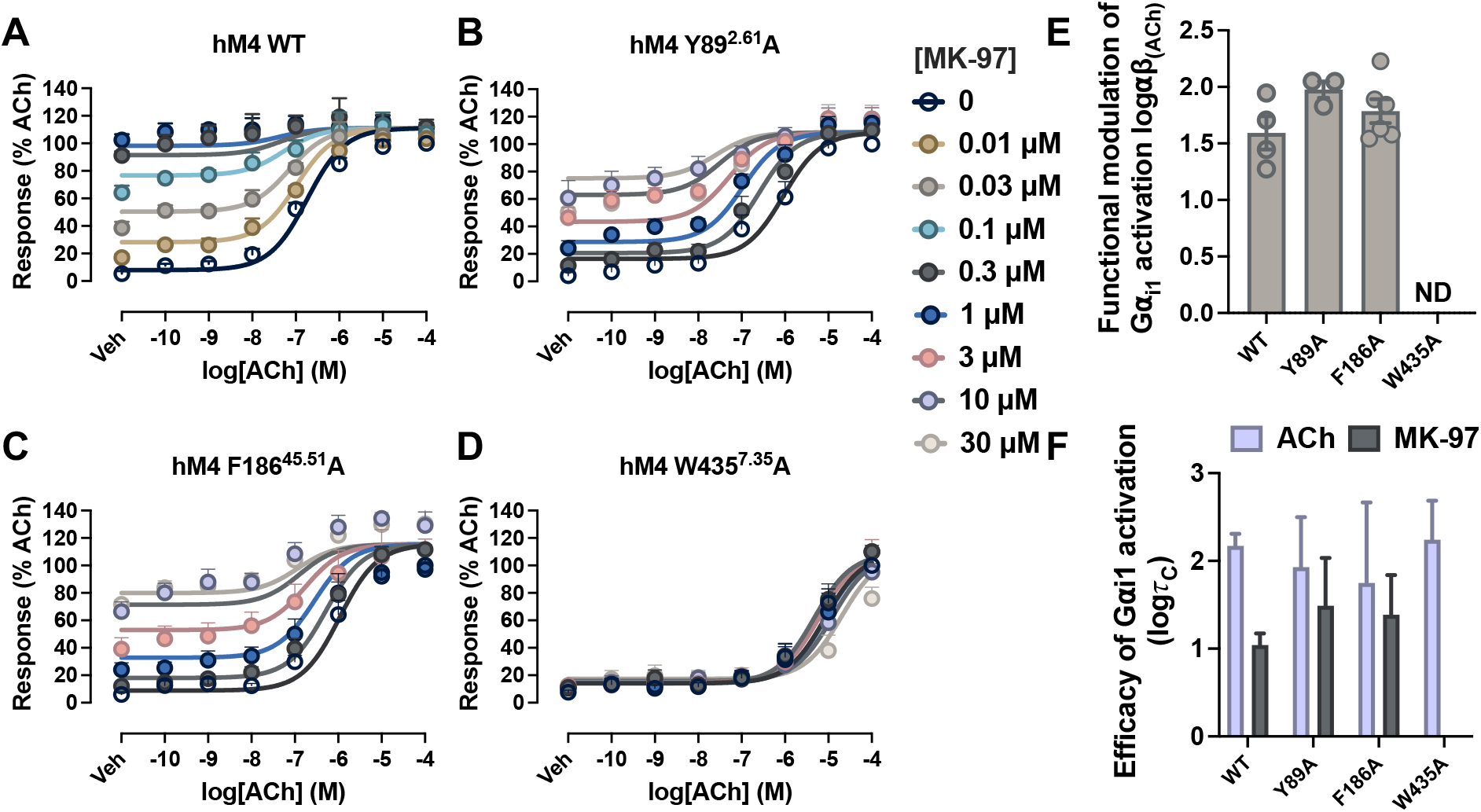
Effect of allosteric site mutations on functional interactions between MK-97 and ACh. (**A-D**) Concentration-response curves of ACh in the presence or absence of MK-97 in Gi1 protein activation at (A) WT and (B-D) mutant M_4_ mAChRs. Data are the mean ± SEM from at least three independent experiments performed in duplicate and normalized to the maximal response of ACh (100%) and vehicle effects (0%). (**E-F**) Pharmacological parameters were estimated by fitting an operational model of allosterism and agonism to estimate the (E) functional allosteric modulation (α*β*) between ACh and MK-97 and (F) logarithm of the operational efficacy parameter corrected for receptor expression (*τ*_C_). ND: value not determined.

Mutations Y89A^2.61^ and F186A^45.51^ had no significant effect on the functional modulation (*αβ*) imparted by MK-97, nor on the signaling efficacy (*τ*) of either ACh or MK-97 after correction for differences in receptor expression levels (**Figure 5E,F**). These findings indicate that while Y89^2.61^ and F186^45.51^ contribute to MK-97 binding affinity, they do not play critical roles in receptor-mediated G protein activation. In contrast, mutation of W435A^7.35^ completely abolished both the allosteric agonism of MK-97 and the functional cooperativity between the PAM and ACh (**Figure 5D,F**). This complete loss of allosteric function is consistent with the critical role of W435^7.35^ in MK-97 binding affinity and cooperativity observed in our binding studies, reinforcing that this residue serves as an essential anchor point for both PAM engagement and allosteric signal transmission.

### MD Simulations

To evaluate the stability of MK-97 in the allosteric site, we performed Gaussian accelerated Molecular Dynamics (GaMD) simulations on the M_4_R-G_i1_-ACh-MK-97 complex. Six independent 500 ns GaMD production simulations showed that ACh remained stable in the orthosteric site, with ~1-2 Å RMSDs relative to the cryo-EM structure (**Figure 6A,C**). For MK-97, the cryo-EM structure represented the lowest energy bound conformation, although it could sample an additional minor intermediate conformation with the GaMD boost potential in two of the six GaMD simulations (Sim1 and Sim2 in **Figure 6B-D)**. Meanwhile, MK-97 formed a π-π stacking interactions with W435^7.35^ in the allosteric site despite fluctuations in the χ_2_ dihedral angle of W435^7.35^ and their interacting distance (**Figure 6E,F**). The key interactions observed in the cryo-EM structure were maintained throughout most of the GaMD simulations, other than Sim1 and Sim2, with Y89^2.61^ and Y92^2.64^ forming hydrogen bonds with MK-97 (**Figure 6G,H**) and the π-π stacking interaction between W435^7.35^ and MK-97, as noted above (**Figure 6F**). The simulations also revealed that F186^45.51^ formed rather weak π-π interactions with the isoindolinone pendant of MK-97 (**Figure 6I**), consistent with the pharmacological observation that the F186A^45.51^ mutation reduces but does not abolish MK-97 binding affinity or functional modulation, unlike the complete loss of activity observed for first-generation PAMs at this mutant^22^. The toggle switch W413^6.48^ underwent only slight fluctuations, adopting mostly 120° and 60° in the χ_2_ dihedral angle of its side chain **(Figure 6J)**. The distance between R130^3.50^ - T399^6.34^ remained stable at ~13-16 Å and the center-of-mass distance between the receptor NPxxY motif and the Gα C-terminus stayed at ~12-13 Å across all simulations, collectively indicating that MK-97 binding maintains the active-state receptor-G protein interface (**Figure 6K,L**). The GaMD simulations validate the structural observations and demonstrate that the distributed interaction network between the M_4_ mAChR bound to MK-97 in the absence and presence of orthosteric ligand facilitates multiple complementary contacts, rather than dependence on minimal critical interactions.

**Figure 6:**
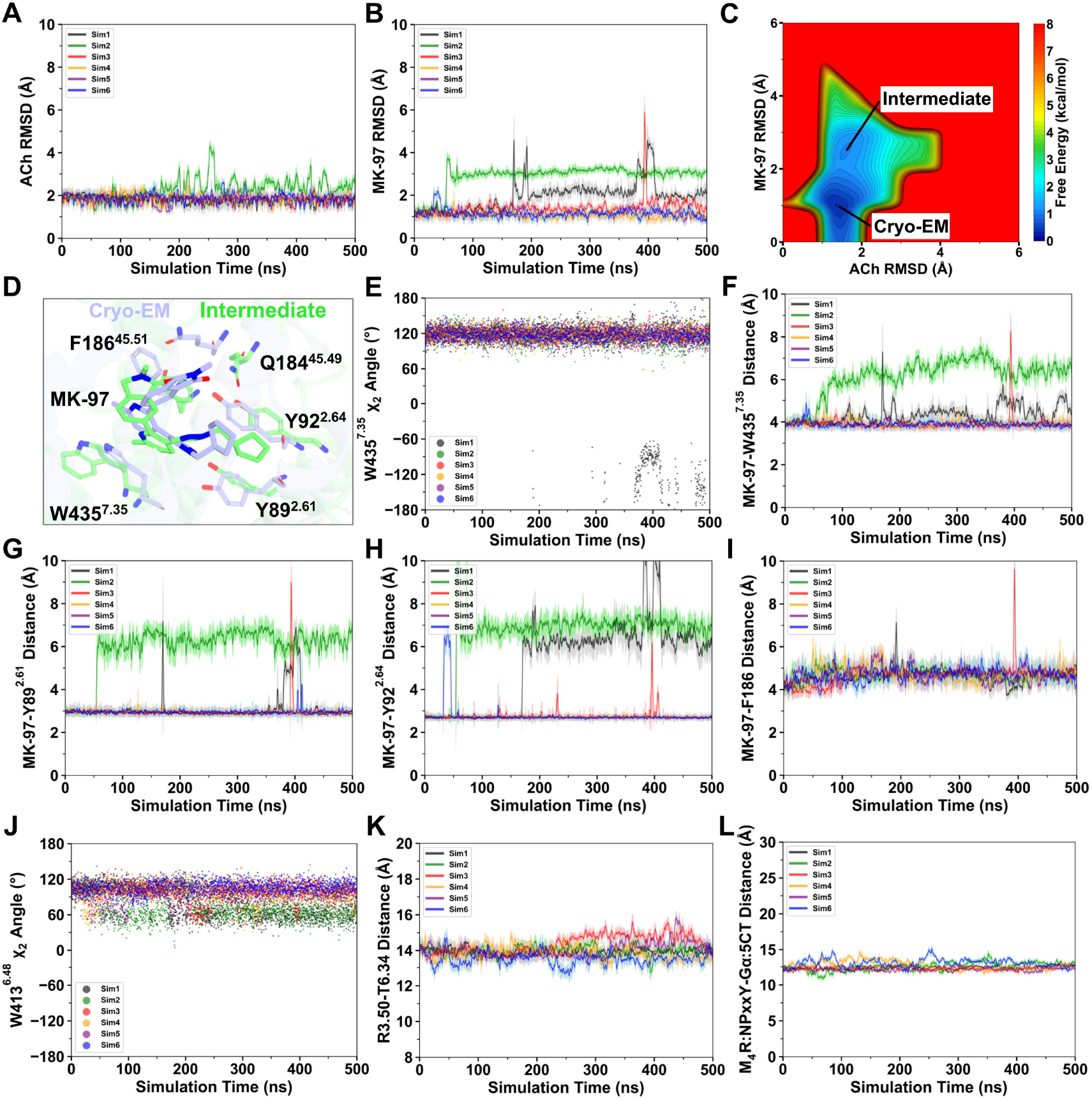
GaMD simulations of the M_4_R-G_i1_-ACh-MK-97 complex. Time courses of the root mean square deviations (RMSDs) of (**A**) agonist ACh **(B)** and the PAM MK-97 relative to the cryo-EM structure. 2D free energy profile of MK-97 RMSD and ACh RMSD with two low-energy states identified, including the cryo-EM structure as the lowest energy conformation and **(C)** an additional minor intermediate conformation of the PAM. (**D**) Overlay of the “Cryo-EM” and “Intermediate” conformations of MK-97. (**E**) Time courses of the χ_2_ dihedral angle in W435^7.35^ obtained from the GaMD simulations. (**F-I**) Time courses of the center-of-mass (F) distance between the pyridine of MK-97 and indole ring of W435^7.35^, (G) distance between the pyrazole nitrogen atom of MK-97 and hydroxyl oxygen atom of Y89^2.61^, (H) distance between the carbonyl oxygen atom of the isoindolinone in MK-97 and hydroxyl oxygen atom of Y92^2.64^, and (I) center-of-mass distance between the pyridine of MK-97 and phenyl ring of F186^45.51^. (**J**) Time courses of the χ_2_ dihedral angle in the W413^6.48^ toggle switch. (**K**) Time courses of the distance between the Cα atoms of R130^3.50^ and T399^6.34^ and (**L**) center-of-mass distance between the NPxxY motif in M_4_R and the last five C-terminal residues in the α5 helix of Gα.

## Discussion

The development of M_4_ mAChR-selective PAMs has historically been hindered by relatively intractable structure-activity relationships, species variability, and poor pharmacokinetic properties^35,36,43,58,59^. These challenges have prevented clinical translation and resulted in limited structural diversity among M_4_ mAChR PAM scaffolds. Here, we report the first cryo-EM structure of a second-generation pyrazol-4-ylpyridine-based M_4_ mAChR PAM, providing critical insights into the molecular basis for the enhanced affinity and distinct pharmacological properties that distinguish MK-97 from first-generation mAChR allosteric modulators.

Our 2.7 Å structure reveals that MK-97 adopts a ‘boomerang’-shaped conformation in the ECV, engaging the allosteric site through a distributed interaction network across three distinct regions. This extended binding mode differs from the planar poses of LY2033298 and VU0467154, potentially explaining the reduced susceptibility of MK-97 to single-residue mutations and approximately 10-fold greater affinity for the allosteric site even in the absence of ACh. Our M_4_R-MK-97 structure helps rationalize extensive structure-activity relationships (SAR) reported for pyrazol-4-ylpyridine analogs^42,43^, revealing why specific structural modifications enhance or diminish M_4_ mAChR PAM activity.

The cyclopentylmethyl substituent on the pyrazole ring, which forms the lower arm of the MK-97 boomerang conformation, occupies a deep hydrophobic pocket formed by Y89^2.61^, Y92^2.64^, I93^2.65^, F186^45.51^, and W435^7.35^. This structural observation explains the SAR finding that the cyclopentylmethyl group provides optimal affinity and functional properties among *N*-alkyl substituents tested^42,43^. Liu *et al*. demonstrated that replacing this group with polar or charged functionalities (carboxylic acids, esters, amides, or amines) caused 10-fold or greater losses in binding affinity, which our structure attributes to the disruption of essential hydrophobic contacts in the allosteric pocket. Similarly, the observation that branched tertiary amine substitution (compound **13** in *Liu et al*.; **Figure 7**) maintained allosteric properties despite reduced affinity suggests this modification can be accommodated within the pocket, albeit with suboptimal packing. The complete loss of activity observed when the *N*-cyclopentylmethyl group was replaced with shorter methylene-linked substituents reflects insufficient reach of the PAM into the deep hydrophobic pocket, likely preventing proper stabilization of the pyrazole-Y89^2.61^ hydrogen bond that our mutagenesis studies confirmed was critical for PAM binding affinity.

**Figure 7:**
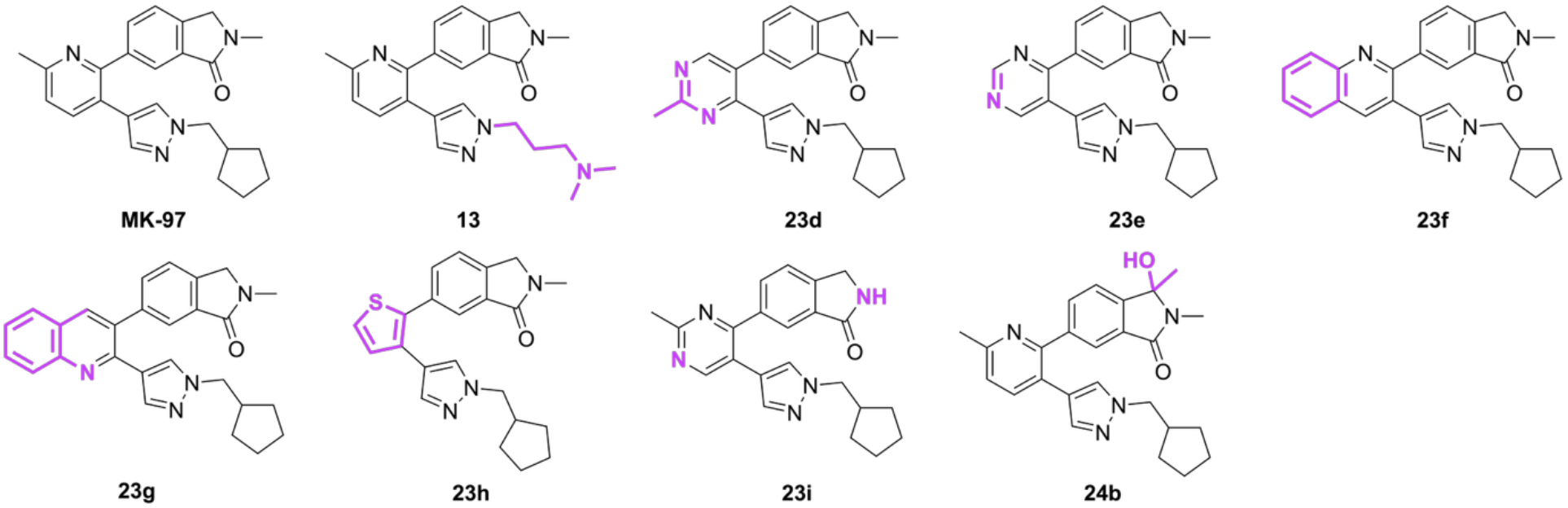
Structural analogs of MK-97. Regions of difference to MK-97 are colored purple. Compounds are from prior publications.^42,43^

The central pyridine vertex serves as the scaffold anchor, forming a π-π interaction with W435^7.35^ that is essential for M_4_ mAChR PAM activity reported to date within the ECV site. Our structure reveals why modifications to this core PAM region produce varied pharmacological profiles. Both Liu *et al*. and Jörg *et al*. explored alternative heterocycles including pyrimidines (compounds **23d, 23e**), quinolines (**23f, 23g**), and a thiophene (**23h**) (**Figure 7**), observing divergent effects on affinity, cooperativity, and efficacy^42,43^. The reduced affinity of most PAM core replacements can be attributed to altered geometry that disrupts the precise positioning required for simultaneous engagement of all three pendant arms.

Notably, the quinoline analog **23f**^42,43^ maintained similar affinity to MK-97 while displaying enhanced allosteric modulation in cAMP signaling, suggesting that the extended aromatic system can engage additional interactions with the allosteric site. In contrast, the quinoline regioisomer **23g** was poorly tolerated (**Figure 7**). This dramatic difference suggests that the nitrogen position within the quinoline core critically influences the electronic properties and conformational preferences of the central scaffold, altering the quality of the π-π stacking interaction with W435^7.35^ or the overall dipole moment of the molecule. This electronic constraint suggests that successful core modifications must preserve not only appropriate geometry for π-stacking, but also the specific electronic character required for optimal allosteric site engagement. The pyrimidine analogs **23d** and **23e** further support this principle, as these compounds retained affinity but became neutral allosteric ligands with minimal cooperativity or efficacy. That is, the pyridine nitrogen and electronic properties appear to optimize not just binding affinity, but also the ability to propagate allosteric effects to the orthosteric site and G protein coupling.

The isoindolinone moiety at the upper arm extends toward ECL2, forming interactions that appear unique to second-generation PAMs disclosed to date. Our structure shows that the isoindolinone carbonyl makes a hydrogen bond with Y92^2.64^, while the *N*-methyl group is oriented towards Q184^45.49^ and the solvent-exposed extracellular region. This structural insight explains the SAR observations that modifications to the isoindolinone had minimal effects on most pharmacological parameters^42,43^. The orientation of the *N*-methyl group toward Q184^45.49^ and the solvent-exposed extracellular region, rather than a constrained hydrophobic pocket, accounts for the tolerance observed with the desmethyl analog **23i**, which maintained allosteric properties and efficacy. Notably, the Y92^2.64^A mutation had minimal effect on MK-97 activity despite the observed hydrogen bond between the isoindolinone carbonyl and Y92^2.64^, suggesting that this interaction is not essential for binding or function and further explaining why this region tolerates structural modifications. The preservation of activity with 3-hydroxy substitution on the isoindolinone (compound **24b** in Liu *et al*.) is consistent with this position being solvent-exposed and able to accommodate additional polar functionality, presenting opportunities for elaboration to improve pharmacokinetic properties or introduce favorable interactions with Q184^45.49^ or neighboring ECL2 residues in next-generation analogs.

Beyond rationalizing existing SAR, our cryo-EM structure provides insights into potential mechanisms underlying the improved mutational tolerance of MK-97 compared to first-generation PAMs. While multiple single-residue mutations, including F186^45.51^A and Y89^2.61^A, completely abolish first-generation PAM activity^22,45,52^, MK-97 retains measurable binding and cooperativity at these mutants. Additionally, the Y92^2.64^A mutation had minimal effects on MK-97 despite an observed hydrogen bond with the isoindolinone carbonyl. These observations are consistent with a distributed interaction network spanning three distinct structural regions.

## Conclusions

The structural and mechanistic insights reported here establish a framework for the rational design of improved M_4_ mAChR-targeted therapeutics. The MK-97 binding mode defines a structural template for next-generation PAM optimisation, while the comprehensive SAR understanding and mutagenesis validation presented here provide critical guidance for the development of M_4_ mAChR therapeutics with improved selectivity, reduced species variability, and enhanced clinical potential for the treatment of neurological disorders.

## Experimental Section

### Synthesis of MK-97

MK-97 was synthesized and characterized as previously described^42,43^. Compound purity was assessed by NMR, LC-MS, and HRMS data that confirmed >95% purity (**Supporting Information**).

### Cell Culture

*Spodoptera frugiperda* (Sf9) and *Trichoplusia ni* (Tni) cells (Expression Systems, Davis, CA, USA) were maintained in ESF-921 media (Expression Systems) at 27 °C while shaking. Flp-In Chinese Hamster Ovary (CHO) cells (Thermo Fisher Scientific) stably expressing the human WT or mutant M_4_ mAChRs were grown in Dulbecco’s Modified Eagle’s Medium (DMEM, Invitrogen) containing 5% FBS and 600 μg/mL hygromycin in a humidified incubator at 37 °C (5% CO_2_, 95% O_2_).

#### Radioligand Binding Assays

Flp-In CHO cells stably expressing M_4_ mAChR constructs were seeded at 1.0 × 10^4^ cells/well and 5.0 × 10^4^ cells/well for WT and mutant receptors, respectively, in 96-well white clear bottom isoplates (Greiner Bio-one) and allowed to adhere overnight in an incubator (37 °C, 5% CO_2_, 95% O_2_). Saturation binding assays were performed to quantify receptor expression and equilibrium dissociation constants with the radioligand [^3^H]-scopolamine methyl chloride ([^3^H]-NMS) (specific activity, 70 – 80 Ci/mmol; PerkinElmer), or [^3^H]-L-quinuclidinyl benzilate (QNB) (specific activity, 250 μCi; PerkinElmer) as previously described^17^. The plates were washed with phosphate-buffered saline (PBS) and cells were incubated with 10 μM of atropine and 0.01 – 10 nM of radioactive ligand in binding buffer, 1× Hank’s Balanced Salt Solution, 10 mM HEPES pH 7.4, at a final volume of 100 μL, followed by 18 h incubation at RT.

Full interaction radioligand binding assays were performed by incubating Flp-In CHO cells stably expressing M_4_ mAChR constructs with a fixed (p*K*_D_) concentration of [^3^H]-NMS, determined from saturation assays, and varying concentrations of the non-radiolabeled orthosteric agonist, ACh, in the absence or presence of increasing concentrations of the allosteric modulator, MK-97, in binding buffer. Vehicle effects were determined with 0.1% dimethylsulfoxide (DMSO) and the reaction was left to reach equilibrium for 18 h at RT. In all cases, the assays were terminated the next day by washing the plates twice with cold 0.9% NaCl to remove unbound radioligand. Radioactivity was measured using a MicroBeta radiometric detector (PerkinElmer).

#### G Protein Activation Assay

Flp-In CHO cells stably expressing M_4_ mAChR constructs were seeded at a density of 20,000 cells/well in 96-well CulturPlates (PerkinElmer) and allowed to adhere for 4 h at 37 °C, 5% CO_2_, and 95% O_2_. The cells were then transiently transfected using polyethylenimine (PEI, Polysciences) and 20 ng per well of each Gα_i1_-RLuc8, Gβ_3_, Gγ_9_-GFP2 TruPath (Addgene) construct, at a ratio of 1:1:1 with 60 ng of total DNA per well, and further incubated for 48 h. The cells were washed twice with PBS and replaced with 70 μL 1× Hanks Balanced Salt Solution pH 7.4, followed by 30 min incubation at 37 °C. 10 μL of 1.3 μM Prolume Purple coelenterazine (Nanolight Technologies) was added to each well and incubated for 5 – 10 min before bioluminescence resonance energy transfer (BRET) measurements were detected on a PHERAstar plate reader (BMG Labtech) using 410/80 nm and 515/30 nm filters. Baseline measurements were taken for 2 cycles before addition of drugs or vehicle to give a final assay volume of 100 μL and further reading for 10 min. Measurement data were collected and expressed as the BRET ratio of 515/ 30 Gγ_y_-GFP2 emission over 410/80 Gα_i1_-RLuc8 emission. The ratio was corrected using the initial 2 cycle baseline read and then baseline corrected again using the vehicle-treated wells. Data were then normalized using the maximal agonist response of ACh to allow for grouping of results using an area under the curve analysis in Prism. Protein Expression

The human M_4_ mAChR gene was modified to exclude residues 242 – 387 of ICL3 and the N-terminal glycosylation sites were mutated to aspartate (N3D, N9D, and N13D) (**Supplementary Figure 1A**) ^22^. This construct and 6× histidine (HIS)-tagged Gβ_1_γ_2_ subunits were cloned into a pVL1392 vector. DNGα_i1_ was cloned into a pFastBac vector^22^. Receptor was expressed in *Spodoptera frugiperda* (Sf9) cells and the DNGα_i1_Gβ_1_γ_2_ complex was expressed in *Trichoplusia ni* (Tni) cells. Cells were grown in ESF-921 and infected at a density of 4.0 × 10^6^ cells per mL, at a ratio of 1:1 for co-infection with DNGα_i1_ and Gβ_1_γ_2_. Expression of the M_4_ mAChR was supplemented with 10 µM atropine. Cells were grown at 27 °C for 60 h post infection and harvested by centrifugation (5000 × g, 15 min, 4 °C)^22^. Pellets were stored at –80 °C.

#### Protein Purification

Purification of the protein complex was as previously described^22^. Tni insect cells were infected with scFv16 baculovirus at a density of 4.0 × 10^6^ cells per mL and incubated for 60 h, followed by harvest by centrifugation (10,000 × g, 10 min, 4 °C). The supernatant was pH balanced to 7.5 and 5 mM CaCl_2_ was added to quench any chelating agents and left to stir for 1.5 h at RT. The supernatant was centrifuged (30,000 × g, 15 min) to remove any precipitants before the addition of 5 mL of EDTA-resistant Ni resin (Cytiva, Marlborough, MA, USA). After 2 h incubation at 4 °C while stirring, the resin was collected in a glass column and washed with 20 column volumes (CVs) of high salt buffer (20 mM HEPES pH 7.5, 100 mM NaCl, 20 mM imidazole) followed by 20 CVs of low salt buffer (20 mM HEPES pH 7.5, 100 mM NaCl, 20 mM imidazole). Protein was eluted using 8 CV of elution buffer (20 mM HEPES pH 7.5, 100 mM NaCl, 250 mM imidazole) and concentrated using a 100 kDa Amicon filter (Millipore, Burlington, MA, USA).

Insect cell pellets expressing M_4_ mAChR from 3 L culture was thawed at RT and solubilized for 2 h at 4 °C in solubilization buffer 20 mM HEPES pH 7.5, 750 mM NaCl, 5 mM MgCl_2_, 5 mM CaCl_2_, 10% glycerol, 10 μM atropine, 0.5 % lauryl maltose neopentyl glycol (LMNG; Anatrace, Maumee, OH, USA), 0.02 % cholesterol hemisuccinate (CHS; Antrace), and cOmplete Protease Inhibitor Cocktail (Roche). The solution was dounced until homogeneous and centrifuged at 30 000 × g for 30 min to remove insoluble material before loading onto an M1 anti-Flag affinity resin, which had been equilibrated with high salt buffer. After 1 h, the resin was washed using a peristaltic pump for 30 min at 2 mL/min with high salt buffer, 20 mM HEPES pH 7.5, 750 mM NaCl, 5 mM MgCl_2_, 5 mM CaCl_2_, 0.5% LMNG, and 0.02% CHS followed by a wash with low salt buffer, 20 mM HEPES pH 7.5, 100 mM NaCl, 5 mM MgCl_2_, 5 mM CaCl_2_, 0.5% LMNG, 0.02% CHS, supplemented with 1 mM ACh and 30 μM MK-97. During this time, the DNGα_i1_–Gβ_1_γ_2_ pellet was thawed, solubilized, and dounced in solubilization buffer, 20 mM HEPES pH 7.5, 100 mM NaCl, 5 mM MgCl_2_, 5 mM CaCl_2_, 0.25% LMNG, 0.01% CHS, cOmplete Protease Inhibitor Cocktail, and apyrase (five units) as previously described. The material was centrifuged at 30,000 × g for 30 min and supernatant was filtered through a glass fiber filter (Millipore, Burlington, MA, USA). The solubilized G proteins were added to the immobilized M_4_ mAChR in addition to 2 mM ACh, 10 μM MK-97, 3 mg scFv16, and apyrase (five units) and incubated for 1 h at RT. The anti-Flag resin was then loaded onto a glass column and washed with approximately 20 CVs of low salt washing buffer supplemented with 0.01% LMNG, 0.001% CHS, 2 mM ACh, and 10 μM MK-97. The complex was eluted with size-exclusion chromatography (SEC) buffer, 20 mM HEPES pH 7.5, 100 mM NaCl, 5 mM MgCl_2_, 0.005% LMNG, 0.0005% CHS, 2 mM ACh, and 10 μM MK-97, supplemented with 10 mM EGTA and 0.1 mg/mL FLAG peptide, and concentrated using a 100 kDa Amicon filter (Millipore, Burlington, MA, USA) to a final volume of ≤ 500 μL. The sample was then purified by SEC using the Superdex 200 increase 10/300 column (Cytiva, Marlborough, MA, USA) with SEC buffer. Eluted fractions containing the complex were collected and incubated with 2 mM ACh and 10 μM MK-97 for 30 min on ice, followed by concentrating with an Amicon Ultra-15 100 kDa molecular mass cut-off centrifugal filter unit (Millipore, Burlington, MA, USA) to 8.49 mg/mL and flash frozen with liquid nitrogen for storage at –80 °C (**Supplementary Figure 1B**).

### SDS-PAGE and Coomassie Staining

Purified samples were diluted in 4× sodium dodecyl sulfate (SDS) loading buffer (200 mM Tris-HCl pH 6.8, 400 mM dithiothreitol, 8% SDS, 0.4% bromophenol blue, 40% glycerol) and separated on 10% Mini-PROTEAN TGX Precast Gels (Bio-Rad, Hercules, CA, US) in SDS running buffer (25 mM Tris, 200 mM glycine, 25 mM SDS) at 150 V for 50 min. The gel was then stained with InstantBlue Coomassie Protein Stain (Bio-Rad) (**Supplementary Figure 1C**).

### Cryo-EM Preparation and Data Collection

3 μL of the sample was applied to a glow-discharged UltrAuFoil R1.2/1.3 300 Holey mesh grids (Quantifoil, Großlöbichau, Germany) and vitrified on a Vitrobot Mark IV (Thermo Fisher Scientific) set at 100% humidity and 4 °C. Data were collected on a Titan Krios G3i 300 kV electron microscope (Thermo Fisher Scientific) equipped with GIF Quantum energy filter and K3 detector (Gatan, Pleasanton, CA, USA). The Krios was operated at an accelerating voltage of 300 kV with a 50 *μ*m C2 aperture, 100 μm objective aperture inserted, and zero-loss filtering (25 eV slit width), at 105kX magnification in nanoprobe EFTEM mode corresponding to a pixel size of 0.83 Å. Each video comprised of 71 frames with a total dose of 53.6 e^−^/Å^2^.

### Cryo-EM Image Processing

6129 micrographs were motion corrected using RELION v3.1 implementation of MotionCor2^60,61^ and contrast transfer function (CTF) estimated through CTFFIND version 4.1_62_. Particles were picked from motion corrected micrographs using a trained model on crYOLO^63^. 5,967,579 particles were initially extracted for reference-free 2D classification and used to pick particles for 3D model generation, and 3D refinements. Particles within bad classes were removed and the remaining 413,103 particles were subjected to Bayesian polishing for 3D classification and further 3D refinements^61^. A final set of 347,823 particles underwent CTF refinement and a final 3D refinement in RELION v3.1, which yielded a 2.7 Å (FSC = 0.143, gold standard) resolution map. To improve receptor map quality, the particles were imported into CryoSPARC^64^ and a local refinement was performed using a receptor mask, which yielded a 2.55 Å map (FSC = 0.143, gold standard, **Supplementary Figure 4**).

### Model Building and Refinement

An initial template model of M_4_ mAChR-DNGα_i1_β_1_γ_2_-scFv16 was generated from the previously determined structure of the M_4_ mAChR (PDB:7TRP)^22^. Models were fitted into EM maps using UCSF ChimeraX^65^ and rigid body fit using PHENIX^66^, followed by iterative rounds of model building in COOT^67^ and real-space refinement in PHENIX^66^. Ligand restraints were generated from the GRADE server (https://grade.globalphasing.org). Model validation was performed with MolProbity^68^ and the wwPDB validation server^69^. Figures were generated with ChimeraX. Fix of the model to the cryo-EM maps is shown in **Supplementary Figure 5**.

## Data Analysis

All pharmacological data were analyzed using GraphPad Prism (versions 9.5.1, 10.1.1, 10.4.1). For radioligand saturation binding, non-specific and total binding data were fitted to a one-site binding model, as previously described^70^ to determine receptor density and radioligand affinity. Interaction experiments between [^3^H]-NMS, ACh, and MK-97 were analyzed using an allosteric ternary complex model to estimate binding affinity values for each ligand and the degree of affinity modulation between agonist and PAM:

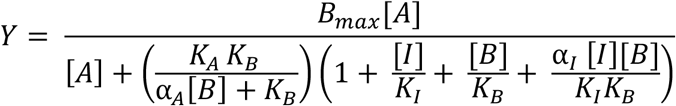

Where Y is the specific radioligand binding, B_max_ is the total number of receptors, [A], [B], and [I] are the concentrations of radioligand, allosteric ligand, and unlabeled orthosteric ligand, respectively. *K*_A_, *K*_B_, and *K*_I_ are the equilibrium dissociation constants of the radioligand, allosteric modulator, and unlabeled orthosteric ligand, respectively, with *K*_A_ fixed to the value determined from saturation binding experiments. The affinity cooperativity values between the allosteric ligand and the radioligand or orthosteric ligand are represented by α_A_ and α_I_, respectively. The global *K*_B_ and α values were determined from grouping each respective value from independent experiments.

For the TruPath assays, data were analyzed using an operational model of allosterism and agonism to determine values of orthosteric (*τ*_A_) and allosteric (*τ*_B_) ligand efficacy and the overall functional modulation (α*β*) between ACh and MK-97^34,54,57^:

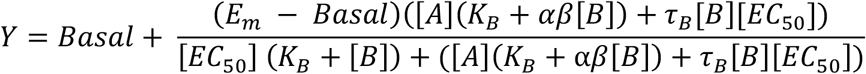

Where E_m_ is the maximal response of the system, Basal is the basal level of response in the absence of agonist, [A] and [B] are the concentrations of the agonist and PAM, respectively. *K*_B_ denotes the equilibrium dissociation constant of the PAM and *τ*_B_ is an operational index of the coupling efficiency/efficacy of the PAM. α represents the binding cooperativity between the orthosteric and allosteric ligands, whereas *β* denotes the allosteric effect of the PAM on the orthosteric ligand efficacy. [EC_50_] denotes the midpoint potency concentration of the orthosteric agonist, A, in the absence of allosteric modulator. The allosteric modulator binding affinity was constrained to the value determined from radioligand binding experiments. The global *τ*_A_, *τ*_B_, and α*β* values were determined from grouping each respective value from independent experiments. For comparison between WT M_4_ mAChR and other M_4_ mAChR constructs, the τ values were corrected (denoted as *τ*_C_) by normalizing value to the B_max_ value for each respective construct determined in saturation binding experiments. This equation is only valid when the allosteric ligand does not change the maximal system response^71^, which was the case for all functional experiments in this study.

All affinity, efficacy, potency and cooperativity values were estimated as logarithms, and statistical analysis between WT and mutant M_4_ mAChRs were determined by one-way ANOVA using a Dunnett’s post-hoc test with a value of P < 0.05 considered as significant in this study.

### Gaussian accelerated Molecular Dynamics (GaMD)

GaMD enhances the conformational sampling of biomolecules by adding a harmonic boost potential to smooth the system potential energy surface^72^. Consider a system with *N* atoms at positions 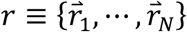. When the system potential 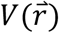 is lower than the threshold energy *E*, a boost potential is added as:

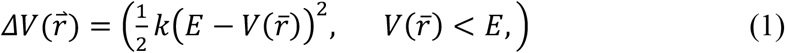

where *k* is the harmonic force constant. The two adjustable parameters E and k are automatically determined on three enhanced sampling principles. First, for any two arbitrary potential values 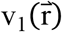 and 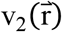 found on the original energy surface, if 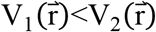, Δ*V* should be a monotonic function that does not change the relative order of the biased potential values; i.e., 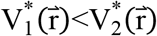. Second, if 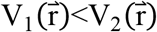, the potential difference observed on the smoothened energy surface should be smaller than that of the original; i.e., 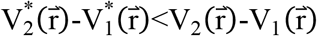. By combining the first two criteria and plugging in the formula of 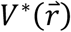 and Δ*V*, we obtain:

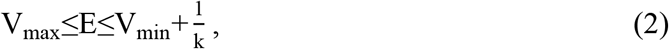

Where V_min_ and V_max_ are the system minimum and maximum potential energies. To ensure that Eq. 2 is valid, *k* has to satisfy: k≤1/(V_max_-V_min_). Let us define: k=k_0_·1/(V_max_-V_min_), then 0<k_0_≤1. Third, the standard deviation (SD) of Δ*V* needs to be small enough (i.e. narrow distribution) to ensure accurate reweighting using cumulant expansion to the second order: σ_ΔV_=kRE-V_avg_Tσ_V_≤σ_0_, where V_avg_ and σ_V_ are the average and SD of ΔVwith σ_0_ as a user-specified upper limit (e.g., 10k_B_T) for accurate reweighting. When E is set to the lower bound E=V_max_ according to Eq. 2, k_0_ can be calculated as:

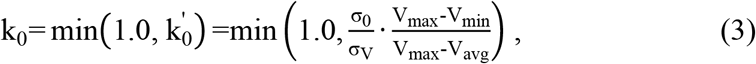

Alternatively, when the threshold energy E is set to its upper bound E=V_min_+1/k, k_0_ is set to:

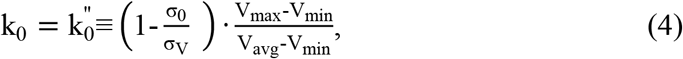

If 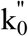 is calculated between 0 and 1. Otherwise, k_0_ is calculated using Eq. 3.

### GaMD simulations and Analysis

Cryo-EM structure of the M_4_ mAChR in complex with its cognate Gα_i1_ heterotrimer, co-bound to ACh and MK-97, were used to set up the GaMD simulation system. The intracellular loop 3 of the receptor and the α-helical domain of the G-protein, which were missing in the cryo-EM structure, were not modelled. The simulation system was prepared by inserting the ACh-MK97-bound M_4_ mAChR-Gα_i1_ complex into a POPC (palmitoyl-2-oleoyl-sn-glycero 3-phosphocholine) lipid bilayer using the CHARMM-GUI online server^73^. All protein chain termini were capped with Acetyl and N-methylamide neutral groups and the disulfide bonds (C105-C185 and C426-C429) were maintained in the M_4_ mAChR. The protein and lipid bilayer were solvated with TIP3P water molecules in a box of 90.0Å × 90.0Å × 150.0Å with the periodic boundary condition. The system charge was neutralized with 0.15 M NaCl. The AMBER FF19SB force field was applied for the protein and LIPID21 for the lipids^74,75^. The general amber force field with GAFF2 parameters were used for ACh and MK-97^76^.

The simulation system was first energy minimized for 5000 steps with constraints on heavy atoms of the proteins and phosphorus atom of the lipids, and a constant number, volume, and temperature (NVT) ensemble equilibration was then performed for 125 ps at 310 K. Using a constant number, pressure, and temperature (NPT) ensemble, additional equilibration was carried out for 375 ps at 310 K. We then performed conventional MD simulation on the system for 20 ns at 1 atm pressure and 310 K temperature with the AMBER22 software package^77^. Long-range electrostatic interactions were computed with the particle mesh Ewald summation method, and a cutoff distance of 9 Å was used for the short-range electrostatic and van der Waals interactions. After cMD, we performed GaMD equilibration for 60 ns, followed by six independent 500-ns GaMD production simulations with randomized initial atomic velocities. All GaMD simulations were performed at the “dual-boost” level, with *α*_0P_, *α*_0D_, and iE parameters set to 12.0 kcal/mol, 12.0 kcal/mol, and 1, respectively. All the six GaMD production trajectories were combined for analysis. The CPPTRAJ software tool was applied to calculate the time-courses of the root-mean-square deviations (RMSDs) of ACh agonist and MK-97 relative to the simulation starting structure, as well as the dihedral angle for W435^7.35 78^. Time courses were also plotted for the hydrogen bond distance between MK-97 and the Y89^2.61^ and Y92^2.64^ residues in M4 mAChR, as well as the π-π stacking interaction between pyridine ring of MK-97 and W435^7.35^ residue in the M_4_ mAChR.

## Associated Content

Supporting Information

PDB ID codes: 23OP

## Acknowledgements

This work was partially supported by the Monash University Ramaciotti Centre for cryo-electron microscopy and the Monash University MASSIVE high-performance computing facility and supercomputing resources with the XSEDE allocation award TG-MCB180049, BIO220137 from the Advanced Cyberinfrastructure Coordination Ecosystem: Services & Support (ACCESS) program, and NERSC project M2874. Molecular graphics and analyses for the cryo-EM and molecular dynamics simulation datasets were produced with Chimera and ChimeraX from the Resource for Biocomputing, Visualization, and Informatics at the University of California, San Francisco.

## Notes

A.C and P.M.S are co-founders, shareholders, and scientific advisors for Septerna Inc. D.W is a scientific advisory board member and shareholder for Septerna Inc. The remaining authors declare no competing financial interest.

## Author Contributions

D.M.T., C.V., A.C., and A.B.T. designed the project. Z.V. purified the M_4_ mAChR-ACh-MK-97 complex. R.D. collected the cryo-EM data. M.J.B. performed the initial cryo-EM processing. M.G.K. and J.I.M. performed data processing with cryoSPARC, and performed model building and structure analysis with D.M.T.. F.X., K.J., and J.W. performed molecular dynamics simulations with supervision from Y.M.. M.G.K. performed molecular pharmacology experiments with assistance from N.B., V.P., M.Y., G.T., and E.T.W.. M.J., B.C., and P.J.S. provided MK-97. P.M.S. and D.W. provided support for cryo-EM data collection and processing. M.G.K. and D.M.T. wrote the initial manuscript draft with review from all authors. D.M.T. and C.V. provided overall project supervision.

## Abbreviations

ACh: acetylcholine
BRET: bioluminescence resonance energy transfer
cryo-EM: cryo-electron microscopy
CHO: Chinese hamster ovary
cAMP: cyclic AMP
CHS: cholesterol hemisuccinate
CTF: contrast transfer function
ECV: extracellular vestibule
ECL: extracellular loop
GPCR: G protein-coupled receptor
GaMD: Gaussian accelerated molecular dynamics
HIS: histidine
ICL: intracellular loop
LMNG: lauryl maltose neopentyl glycol
QNB: L-quinuclidinyl benzilate
mAChR: muscarinic acetylcholine receptor
PAM: positive allosteric modulator
PET: positron emission tomography
PEI: polyethylenimine
RMSD: root mean squared deviation
RT: room temperature
SAR: structure-activity relationship
NMS: scopolamine methyl chloride
SF9: *Spodoptera frugiperda;*
Tni: *Trichoplusia ni*.

## References

(1) Fredriksson, R., Lagerström, M. C., Lundin, L.-G., Schiöth, H. B. The G-Protein-Coupled Receptors in the Human Genome Form Five Main Families. Phylogenetic Analysis, Paralogon Groups, and Fingerprints. Mol Pharmacol 2003, 63 (6), 1256– 1272. 10.1124/mol.63.6.1256.

(2) Hulme, E. C., Birdsall, N. J. M., Buckley, N. J. Muscarinic Receptor Subtypes. Annu. Rev. Pharmacol. Toxicol. 1990, 30 (1), 633–673. 10.1146/annurev.pa.30.040190.003221.

(3) Ince, E., Ciliax, B. J., Levey, A. I. Differential Expression of D1 and D2 Dopamine and M4 Muscarinic Acetylcholine Receptor Proteins in Identified Striatonigral Neurons. Synapse 1997, 27 (4), 357–366. 10.1002/(SICI)1098-2396(199712)27:4%253C357::AID-SYN9%253E3.0.CO;2-B.

(4) Levey, A. I. Immunological Localization of M1–M5 Muscarinic Acetylcholine Receptors in Peripheral Tissues and Brain. Life Sciences 1993, 52 (5), 441–448. 10.1016/0024-3205(93)90300-R.

(5) Bubser, M., Bridges, T. M., Dencker, D., Gould, R. W., Grannan, M., Noetzel, M. J., Lamsal, A., Niswender, C. M., Daniels, J. S., Poslusney, M. S., Melancon, B. J., Tarr, J. C., Byers, F. W., Wess, J., Duggan, M. E., Dunlop, J., Wood, M. W., Brandon, N. J., Wood, M. R., Lindsley, C. W., Conn, P. J., Jones, C. K. Selective Activation of M4 Muscarinic Acetylcholine Receptors Reverses MK-801-Induced Behavioral Impairments and Enhances Associative Learning in Rodents. ACS Chem. Neurosci. 2014, 5 (10), 920–942. 10.1021/cn500128b.

(6) Bubser, M., Byun, N., Wood, M. R., Jones, C. K. Muscarinic Receptor Pharmacology and Circuitry for the Modulation of Cognition. In Muscarinic Receptors; Fryer, A. D., Christopoulos, A., Nathanson, N. M., Eds., Handbook of Experimental Pharmacology; Springer: Berlin, Heidelberg, 2012; pp 121–166. 10.1007/978-3-642-23274-9_7.

(7) Foster, D. J., Wilson, J. M., Remke, D. H., Mahmood, M. S., Uddin, M. J., Wess, J., Patel, S., Marnett, L. J., Niswender, C. M., Jones, C. K., Xiang, Z., Lindsley, C. W., Rook, J. M., Conn, P. J. Antipsychotic-like Effects of M4 Positive Allosteric Modulators Are Mediated by CB2 Receptor-Dependent Inhibition of Dopamine Release. Neuron 2016, 91 (6), 1244–1252. 10.1016/j.neuron.2016.08.017.

(8) Paul, S. M., Yohn, S. E., Popiolek, M., Miller, A. C., Felder, C. C. Muscarinic Acetylcholine Receptor Agonists as Novel Treatments for Schizophrenia. AJP 2022, 179 (9), 611–627. 10.1176/appi.ajp.21101083.

(9) Dencker, D., Wörtwein, G., Weikop, P., Jeon, J., Thomsen, M., Sager, T. N., Mørk, A., Woldbye, D. P. D., Wess, J., Fink-Jensen, A. Involvement of a Subpopulation of Neuronal M4 Muscarinic Acetylcholine Receptors in the Antipsychotic-like Effects of the M1/M4 Preferring Muscarinic Receptor Agonist Xanomeline. J. Neurosci. 2011, 31 (16), 5905–5908. 10.1523/JNEUROSCI.0370-11.2011.

(10) Zhou, F.-M., Wilson, C., Dani, J. A. Muscarinic and Nicotinic Cholinergic Mechanisms in the Mesostriatal Dopamine Systems. Neuroscientist 2003, 9 (1), 23–36. 10.1177/1073858402239588.

(11) Leach, K., Loiacono, R. E., Felder, C. C., McKinzie, D. L., Mogg, A., Shaw, D. B., Sexton, P. M., Christopoulos, A. Molecular Mechanisms of Action and In Vivo Validation of an M4 Muscarinic Acetylcholine Receptor Allosteric Modulator with Potential Antipsychotic Properties. Neuropsychopharmacol 2010, 35 (4), 855–869. 10.1038/npp.2009.194.

(12) Chan, W. Y., McKinzie, D. L., Bose, S., Mitchell, S. N., Witkin, J. M., Thompson, R. C., Christopoulos, A., Lazareno, S., Birdsall, N. J. M., Bymaster, F. P., Felder, C. C. Allosteric Modulation of the Muscarinic M4 Receptor as an Approach to Treating Schizophrenia. Proceedings of the National Academy of Sciences 2008, 105 (31), 10978–10983. 10.1073/pnas.0800567105.

(13) Byun, N. E., Grannan, M., Bubser, M., Barry, R. L., Thompson, A., Rosanelli, J., Gowrishankar, R., Kelm, N. D., Damon, S., Bridges, T. M., Melancon, B. J., Tarr, J. C., Brogan, J. T., Avison, M. J., Deutch, A. Y., Wess, J., Wood, M. R., Lindsley, C. W., Gore, J. C., Conn, P. J., Jones, C. K. Antipsychotic Drug-Like Effects of the Selective M4 Muscarinic Acetylcholine Receptor Positive Allosteric Modulator VU0152100. Neuropsychopharmacol 2014, 39 (7), 1578–1593. 10.1038/npp.2014.2.

(14) Shannon, H. E., Rasmussen, K., Bymaster, F. P., Hart, J. C., Peters, S. C., Swedberg, M. D. B., Jeppesen, L., Sheardown, M. J., Sauerberg, P., Fink-Jensen, A. Xanomeline, an M1/M4 Preferring Muscarinic Cholinergic Receptor Agonist, Produces Antipsychotic-like Activity in Rats and Mice. Schizophrenia Research 2000, 42 (3), 249–259. 10.1016/S0920-9964(99)00138-3.

(15) Cookson, J., Jonsson, F. A New Cholinergic Mechanism for Antipsychotics: Emraclidine and M4 Muscarinic Receptors. The Lancet 2022, 400 (10369), 2159–2161. 10.1016/S0140-6736(22)02421-7.

(16) Smith, C. M., Augustine, M. S., Dorrough, J., Szabo, S. T., Shadaram, S., Hoffman, E. O. G., Muzyk, A. Xanomeline-Trospium (CobenfyTM) for Schizophrenia: A Review of the Literature. Clin Psychopharmacol Neurosci 2025, 23 (1), 2–14. 10.9758/cpn.24.1253.

(17) Thal, D. M., Sun, B., Feng, D., Nawaratne, V., Leach, K., Felder, C. C., Bures, M. G., Evans, D. A., Weis, W. I., Bachhawat, P., Kobilka, T. S., Sexton, P. M., Kobilka, B. K., Christopoulos, A. Crystal Structures of the M1 and M4 Muscarinic Acetylcholine Receptors. Nature 2016, 531 (7594), 335–340. 10.1038/nature17188.

(18) Haga, K., Kruse, A. C., Asada, H., Yurugi-Kobayashi, T., Shiroishi, M., Zhang, C., Weis, W. I., Okada, T., Kobilka, B. K., Haga, T., Kobayashi, T. Structure of the Human M2 Muscarinic Acetylcholine Receptor Bound to an Antagonist. Nature 2012, 482 (7386), 547–551. 10.1038/nature10753.

(19) Vuckovic, Z., Gentry, P. R., Berizzi, A. E., Hirata, K., Varghese, S., Thompson, G., van der Westhuizen, E. T., Burger, W. A. C., Rahmani, R., Valant, C., Langmead, C. J., Lindsley, C. W., Baell, J. B., Tobin, A. B., Sexton, P. M., Christopoulos, A., Thal, D. M. Crystal Structure of the M5 Muscarinic Acetylcholine Receptor. Proceedings of the National Academy of Sciences 2019, 116 (51), 26001–26007. 10.1073/pnas.1914446116.

(20) Maeda, S., Qu, Q., Robertson, M. J., Skiniotis, G., Kobilka, B. K. Structures of the M1 and M2 Muscarinic Acetylcholine Receptor/G-Protein Complexes. Science 2019, 364 (6440), 552–557. 10.1126/science.aaw5188.

(21) Kruse, A. C., Hu, J., Pan, A. C., Arlow, D. H., Rosenbaum, D. M., Rosemond, E., Green, H. F., Liu, T., Chae, P. S., Dror, R. O., Shaw, D. E., Weis, W. I., Wess, J., Kobilka, B. K. Structure and Dynamics of the M3 Muscarinic Acetylcholine Receptor. Nature 2012, 482 (7386), 552–556. 10.1038/nature10867.

(22) Vuckovic, Z., Wang, J., Pham, V., Mobbs, J. I., Belousoff, M. J., Bhattarai, A., Burger, W. A., Thompson, G., Yeasmin, M., Nawaratne, V., Leach, K., van der Westhuizen, E. T., Khajehali, E., Liang, Y.-L., Glukhova, A., Wootten, D., Lindsley, C. W., Tobin, A., Sexton, P., Danev, R., Valant, C., Miao, Y., Christopoulos, A., Thal, D. M. Pharmacological Hallmarks of Allostery at the M4 Muscarinic Receptor Elucidated through Structure and Dynamics. eLife 2023, 12, e83477. 10.7554/eLife.83477.

(23) Thorsen, T. S., Matt, R., Weis, W. I., Kobilka, B. K. Modified T4 Lysozyme Fusion Proteins Facilitate G Protein-Coupled Receptor Crystallogenesis. Structure 2014, 22 (11), 1657–1664. 10.1016/j.str.2014.08.022.

(24) Burger, W. A. C., Mobbs, J. I., Rana, B., Wang, J., Joshi, K., Gentry, P. R., Yeasmin, M., Venugopal, H., Bender, A. M., Lindsley, C. W., Miao, Y., Christopoulos, A., Valant, C., Thal, D. M. Cryo-EM Reveals a New Allosteric Binding Site at the M5 mAChR. bioRxiv February 8, 2025, p 2025.02.05.636602. 10.1101/2025.02.05.636602.

(25) Andersen, M. B., Fink-Jensen, A., Peacock, L., Gerlach, J., Bymaster, F., Lundbæk, J. A., Werge, T. The Muscarinic M1/M4 Receptor Agonist Xanomeline Exhibits Antipsychotic-Like Activity in Cebus Apella Monkeys. Neuropsychopharmacol 2003, 28 (6), 1168–1175. 10.1038/sj.npp.1300151.

(26) Shekhar, A., Potter, W. Z., Lightfoot, J., Lienemann, J., Dubé, S., Mallinckrodt, C., Bymaster, F. P., McKinzie, D. L., Felder, C. C. Selective Muscarinic Receptor Agonist Xanomeline as a Novel Treatment Approach for Schizophrenia. AJP 2008, 165 (8), 1033–1039. 10.1176/appi.ajp.2008.06091591.

(27) Bender, A. M., Jones, C. K., Lindsley, C. W. Classics in Chemical Neuroscience: Xanomeline. ACS Chem. Neurosci. 2017, 8 (3), 435–443. 10.1021/acschemneuro.7b00001.

(28) Powers, A. S., Pham, V., Burger, W. A. C., Thompson, G., Laloudakis, Y., Barnes, N. W., Sexton, P. M., Paul, S. M., Christopoulos, A., Thal, D. M., Felder, C. C., Valant, C., Dror, R. O. Structural Basis of Efficacy-Driven Ligand Selectivity at GPCRs. Nat Chem Biol 2023, 19 (7), 805–814. 10.1038/s41589-022-01247-5.

(29) Kavoussi, R., Miller, A. C., Brannan, S. K., Breier, A. Xanomeline plus Trospium: A Novel Strategy to Enhance pro-Muscarinic Efficacy and Mitigate Peripheral Side Effects. American Society of Clinical Psychopharmacology 2017.

(30) Correll, C. U., Angelov, A. S., Brannan, S. K. Safety and Efficacy of KarXT (Xanomeline–Trospium) in Patients with Schizophrenia: Results from a Phase 3, Randomised, Double-Blind, Placebo-Controlled Trial (EMERGENT-2). Poster Presentation P. 0193. In 35th ECNP Congress; 2022; pp 15–18.

(31) May, L., Leach, K., Sexton, P., Christopoulos, A. Allosteric Modulation of G Protein-Coupled Receptors. Annual review of pharmacology and toxicology 2007, 47, 1–51. 10.1146/annurev.pharmtox.47.120505.105159.

(32) Burger, W. A. C., Sexton, P. M., Christopoulos, A., Thal, D. M. Toward an Understanding of the Structural Basis of Allostery in Muscarinic Acetylcholine Receptors. Journal of General Physiology 2018, 150 (10), 1360–1372. 10.1085/jgp.201711979.

(33) Christopoulos, A. Allosteric Binding Sites on Cell-Surface Receptors: Novel Targets for Drug Discovery. Nat Rev Drug Discov 2002, 1 (3), 198–210. 10.1038/nrd746.

(34) Leach, K., Sexton, P. M., Christopoulos, A. Allosteric GPCR Modulators: Taking Advantage of Permissive Receptor Pharmacology. Trends in Pharmacological Sciences 2007, 28 (8), 382–389. 10.1016/j.tips.2007.06.004.

(35) Suratman, S., Leach, K., Sexton, P., Felder, C., Loiacono, R., Christopoulos, A. Impact of Species Variability and ‘Probe-Dependence’ on the Detection and in Vivo Validation of Allosteric Modulation at the M4 Muscarinic Acetylcholine Receptor. British Journal of Pharmacology 2011, 162 (7), 1659–1670. 10.1111/j.1476-5381.2010.01184.x.

(36) Wood, M. R., Noetzel, M. J., Poslusney, M. S., Melancon, B. J., Tarr, J. C., Lamsal, A., Chang, S., Luscombe, V. B., Weiner, R. L., Cho, H. P., Bubser, M., Jones, C. K., Niswender, C. M., Wood, M. W., Engers, D. W., Brandon, N. J., Duggan, M. E., Conn, P. J., Bridges, T. M., Lindsley, C. W. Challenges in the Development of an M4 PAM in Vivo Tool Compound: The Discovery of VU0467154 and Unexpected DMPK Profiles of Close Analogs. Bioorganic & Medicinal Chemistry Letters 2017, 27 (2), 171–175. 10.1016/j.bmcl.2016.11.086.

(37) Schubert, J. W., Harrison, S. T., Mulhearn, J., Gomez, R., Tynebor, R., Jones, K., Bunda, J., Hanney, B., Wai, J. M.-C., Cox, C., McCauley, J. A., Sanders, J. M., Magliaro, B., O’Brien, J., Pajkovic, N., Huszar Agrapides, S. L., Taylor, A., Gotter, A., Smith, S. M., Uslaner, J., Browne, S., Risso, S., Egbertson, M. Discovery, Optimization, and Biological Characterization of 2,3,6-Trisubstituted Pyridine-Containing M4 Positive Allosteric Modulators. ChemMedChem 2019, 14 (9), 943–951. 10.1002/cmdc.201900088.

(38) Lange, H. S., Vardigan, J. D., Cannon, C. E., Puri, V., Henze, D. A., Uslaner, J. M. Effects of a Novel M4 Muscarinic Positive Allosteric Modulator on Behavior and Cognitive Deficits Relevant to Alzheimer’s Disease and Schizophrenia in Rhesus Monkey. Neuropharmacology 2021, 197, 108754. 10.1016/j.neuropharm.2021.108754.

(39) Tong, L., Li, W., Lo, M. M.-C., Gao, X., Wai, J. M.-C., Rudd, M., Tellers, D., Joshi, A., Zeng, Z., Miller, P., Salinas, C., Riffel, K., Haley, H., Purcell, M., Holahan, M., Gantert, L., Schubert, J. W., Jones, K., Mulhearn, J., Egbertson, M., Meng, Z., Hanney, B., Gomez, R., Harrison, S. T., McQuade, P., Bueters, T., Uslaner, J., Morrow, J., Thomson, F., Kong, J., Liao, J., Selyutin, O., Bao, J., Hastings, N. B., Agrawal, S., Magliaro, B. C., Monsma, F. J., Smith, M. D., Risso, S., Hesk, D., Hostetler, E., Mazzola, R. Discovery of [11C]MK-6884: A Positron Emission Tomography (PET) Imaging Agent for the Study of M4Muscarinic Receptor Positive Allosteric Modulators (PAMs) in Neurodegenerative Diseases. Journal of medicinal chemistry 2020, 63 (5), 2411–2425. 10.1021/acs.jmedchem.9b01406.

(40) Li, W., Wang, Y., Lohith, T. G., Zeng, Z., Tong, L., Mazzola, R., Riffel, K., Miller, P., Purcell, M., Holahan, M., Haley, H., Gantert, L., Hesk, D., Ren, S., Morrow, J., Uslaner, J., Struyk, A., Wai, J. M.-C., Rudd, M. T., Tellers, D. M., McAvoy, T., Bormans, G., Koole, M., Laere, K. V., Serdons, K., Hoon, J. de; Declercq, R., Lepeleire, I. D., Pascual, M. B., Zanotti-Fregonara, P., Yu, M., Arbones, V., Masdeu, J. C., Cheng, A., Hussain, A., Bueters, T., Anderson, M. S., Hostetler, E. D., Basile, A. S. The PET Tracer [11C]MK-6884 Quantifies M4 Muscarinic Receptor in Rhesus Monkeys and Patients with Alzheimer’s Disease. Science Translational Medicine 2022. 10.1126/scitranslmed.abg3684.

(41) Belov, V., Guehl, N. J., Duvvuri, S., Iredale, P., Moon, S.-H., Dhaynaut, M., Chakilam, S., MacDonagh, A. C., Rice, P. A., Yokell, D. L., Renger, J. J., El Fakhri, G., Normandin, M. D. PET Imaging of M4 Muscarinic Acetylcholine Receptors in Rhesus Macaques Using [11C]MK-6884: Quantification with Kinetic Modeling and Receptor Occupancy by CVL-231 (Emraclidine), a Novel Positive Allosteric Modulator. J Cereb Blood Flow Metab 2024, 0271678X241238820. 10.1177/0271678X241238820.

(42) Liu, B., Thompson, G., Jörg, M., Barnes, N., Thal, D. M., Christopoulos, A., Capuano, B., Valant, C., Scammells, P. J. Discovery of 2-Methyl-5-(1H-Pyrazol-4-Yl)Pyridines and Related Heterocycles as Promising M4 mAChR Positive Allosteric Modulators for the Treatment of Neurocognitive Disorders. J. Med. Chem. 2024, 67 (15), 13286–13304. 10.1021/acs.jmedchem.4c01207.

(43) Jörg, M., van der Westhuizen, E. T., Lu, Y., Christopher Choy, K. H., Shackleford, D. M., Khajehali, E., Tobin, A. B., Thal, D. M., Capuano, B., Christopoulos, A., Valant, C., Scammells, P. J. Design, Synthesis and Evaluation of Novel 2-Phenyl-3-(1H-Pyrazol-4-Yl)Pyridine Positive Allosteric Modulators for the M4 mAChR. European Journal of Medicinal Chemistry 2023, 258, 115588. 10.1016/j.ejmech.2023.115588.

(44) Temple, K. J., Long, M. F., Engers, J. L., Watson, K. J., Chang, S., Luscombe, V. B., Rodriguez, A. L., Niswender, C. M., Bridges, T. M., Conn, P. J., Engers, D. W., Lindsley, C. W. Discovery of Structurally Distinct Tricyclic M4 Positive Allosteric Modulator (PAM) Chemotypes. Bioorganic & Medicinal Chemistry Letters 2020, 30 (4), 126811. 10.1016/j.bmcl.2019.126811.

(45) Burger, W. A. C., Pham, V., Vuckovic, Z., Powers, A. S., Mobbs, J. I., Laloudakis, Y., Glukhova, A., Wootten, D., Tobin, A. B., Sexton, P. M., Paul, S. M., Felder, C. C., Danev, R., Dror, R. O., Christopoulos, A., Valant, C., Thal, D. M. Xanomeline Displays Concomitant Orthosteric and Allosteric Binding Modes at the M4 mAChR. Nat Commun 2023, 14 (1), 5440. 10.1038/s41467-023-41199-5.

(46) Zhou, Q., Yang, D., Wu, M., Guo, Y., Guo, W., Zhong, L., Cai, X., Dai, A., Jang, W., Shakhnovich, E. I., Liu, Z.-J., Stevens, R. C., Lambert, N. A., Babu, M. M., Wang, M.-W., Zhao, S. Common Activation Mechanism of Class A GPCRs. eLife 2019, 8, e50279. 10.7554/eLife.50279.

(47) Shi, L., Liapakis, G., Xu, R., Guarnieri, F., Ballesteros, J. A., Javitch, J. A. β2 Adrenergic Receptor Activation: Modulation of the Proline Kink in Transmembrane 6 by a Rotamer Toggle Switch. Journal of Biological Chemistry 2002, 277 (43), 40989– 40996. 10.1074/jbc.M206801200.

(48) Nygaard, R., Frimurer, T. M., Holst, B., Rosenkilde, M. M., Schwartz, T. W. Ligand Binding and Micro-Switches in 7TM Receptor Structures. Trends in Pharmacological Sciences 2009, 30 (5), 249–259. 10.1016/j.tips.2009.02.006.

(49) Schwartz, T. W., Frimurer, T. M., Holst, B., Rosenkilde, M. M., Elling, C. E. Molecular Mechanism of 7TM Receptor Activation—a Global Toggle Switch Model. Annu. Rev. Pharmacol. Toxicol. 2006, 46, 481–519.

(50) Fritze, O., Filipek, S., Kuksa, V., Palczewski, K., Hofmann, K. P., Ernst, O. P. Role of the Conserved NPxxY(x)5,6F Motif in the Rhodopsin Ground State and during Activation. Proceedings of the National Academy of Sciences 2003, 100 (5), 2290– 2295. 10.1073/pnas.0435715100.

(51) Kruse, A. C., Ring, A. M., Manglik, A., Hu, J., Hu, K., Eitel, K., Hübner, H., Pardon, E., Valant, C., Sexton, P. M., Christopoulos, A., Felder, C. C., Gmeiner, P., Steyaert, J., Weis, W. I., Garcia, K. C., Wess, J., Kobilka, B. K. Activation and Allosteric Modulation of a Muscarinic Acetylcholine Receptor. Nature 2013, 504 (7478), 101– 106. 10.1038/nature12735.

(52) Nawaratne, V., Leach, K., Felder, C. C., Sexton, P. M., Christopoulos, A. Structural Determinants of Allosteric Agonism and Modulation at the M4 Muscarinic Acetylcholine Receptor: IDENTIFICATION OF LIGAND-SPECIFIC AND GLOBAL ACTIVATION MECHANISMS *. Journal of Biological Chemistry 2010, 285 (25), 19012–19021. 10.1074/jbc.M110.125096.

(53) Isberg, V., de Graaf, C., Bortolato, A., Cherezov, V., Katritch, V., Marshall, F. H., Mordalski, S., Pin, J.-P., Stevens, R. C., Vriend, G., Gloriam, D. E. Generic GPCR Residue Numbers - Aligning Topology Maps Minding The Gaps. Trends Pharmacol Sci 2015, 36 (1), 22–31. 10.1016/j.tips.2014.11.001.

(54) Leach, K., Davey, A. E., Felder, C. C., Sexton, P. M., Christopoulos, A. The Role of Transmembrane Domain 3 in the Actions of Orthosteric, Allosteric, and Atypical Agonists of the M4 Muscarinic Acetylcholine Receptor. Mol Pharmacol 2011, 79 (5), 855–865. 10.1124/mol.111.070938.

(55) Haider, A., Deng, X., Mastromihalis, O., Pfister, S. K., Jeppesen, T. E., Xiao, Z., Pham, V., Sun, S., Rong, J., Zhao, C., Chen, J., Li, Y., Connors, T. R., Davenport, A. T., Daunais, J. B., Hosseini, V., Ran, W., Christopoulos, A., Wang, L., Valant, C., Liang, S. H. Structure–Activity Relationship of Pyrazol-4-Yl-Pyridine Derivatives and Identification of a Radiofluorinated Probe for Imaging the Muscarinic Acetylcholine Receptor M4. Acta Pharmaceutica Sinica B 2022. 10.1016/j.apsb.2022.07.008.

(56) Olsen, R. H. J., DiBerto, J. F., English, J. G., Glaudin, A. M., Krumm, B. E., Slocum, S. T., Che, T., Gavin, A. C., McCorvy, J. D., Roth, B. L., Strachan, R. T. TRUPATH, an Open-Source Biosensor Platform for Interrogating the GPCR Transducerome. Nat Chem Biol 2020, 16 (8), 841–849. 10.1038/s41589-020-0535-8.

(57) Christopoulos, A., Kenakin, T. G Protein-Coupled Receptor Allosterism and Complexing. Pharmacol Rev 2002, 54 (2), 323–374. 10.1124/pr.54.2.323.

(58) Valant, C., Felder, C. C., Sexton, P. M., Christopoulos, A. Probe Dependence in the Allosteric Modulation of a G Protein-Coupled Receptor: Implications for Detection and Validation of Allosteric Ligand Effects. Mol Pharmacol 2012, 81 (1), 41–52. 10.1124/mol.111.074872.

(59) Liu, B., Christopoulos, A., Thal, D. M., Capuano, B., Valant, C., Scammells, P. J. The Prosperity and Adversity of M4 Muscarinic Acetylcholine Receptor Activators in the Treatment of Neuropsychiatric Disorders. J. Med. Chem. 2025, 68 (8), 7932–7954. 10.1021/acs.jmedchem.5c00678.

(60) Zheng, S. Q., Palovcak, E., Armache, J.-P., Verba, K. A., Cheng, Y., Agard, D. A. MotionCor2: Anisotropic Correction of Beam-Induced Motion for Improved Cryo-Electron Microscopy. Nat Methods 2017, 14 (4), 331–332. 10.1038/nmeth.4193.

(61) Zivanov, J., Nakane, T., Forsberg, B. O., Kimanius, D., Hagen, W. J., Lindahl, E., Scheres, S. H. New Tools for Automated High-Resolution Cryo-EM Structure Determination in RELION-3. eLife 2018, 7, e42166. 10.7554/eLife.42166.

(62) Rohou, A., Grigorieff, N. CTFFIND4: Fast and Accurate Defocus Estimation from Electron Micrographs. Journal of Structural Biology 2015, 192 (2), 216–221. 10.1016/j.jsb.2015.08.008.

(63) Wagner, T., Merino, F., Stabrin, M., Moriya, T., Antoni, C., Apelbaum, A., Hagel, P., Sitsel, O., Raisch, T., Prumbaum, D., Quentin, D., Roderer, D., Tacke, S., Siebolds, B., Schubert, E., Shaikh, T. R., Lill, P., Gatsogiannis, C., Raunser, S. SPHIRE-crYOLO Is a Fast and Accurate Fully Automated Particle Picker for Cryo-EM. Commun Biol 2019, 2 (1), 1–13. 10.1038/s42003-019-0437-z.

(64) Punjani, A., Rubinstein, J. L., Fleet, D. J., Brubaker, M. A. cryoSPARC: Algorithms for Rapid Unsupervised Cryo-EM Structure Determination. Nat Methods 2017, 14 (3), 290– 296. 10.1038/nmeth.4169.

(65) Meng, E. C., Goddard, T. D., Pettersen, E. F., Couch, G. S., Pearson, Z. J., Morris, J. H., Ferrin, T. E. UCSF ChimeraX: Tools for Structure Building and Analysis. Protein Science 2023, 32 (11), e4792. 10.1002/pro.4792.

(66) Liebschner, D., Afonine, P. V., Baker, M. L., Bunkóczi, G., Chen, V. B., Croll, T. I., Hintze, B., Hung, L. W., Jain, S., McCoy, A. J., Moriarty, N. W., Oeffner, R. D., Poon, B. K., Prisant, M. G., Read, R. J., Richardson, J. S., Richardson, D. C., Sammito, M. D., Sobolev, O. V., Stockwell, D. H., Terwilliger, T. C., Urzhumtsev, A. G., Videau, L. L., Williams, C. J., Adams, P. D. Macromolecular Structure Determination Using X-Rays, Neutrons and Electrons: Recent Developments in Phenix. Acta Crystallogr D Struct Biol 2019, 75 (Pt 10), 861–877. 10.1107/S2059798319011471.

(67) Lohkamp, B., Scott, W. G., Cowtan, K. Features and Development of Coot. Acta Crystallographica. Section D: Biological Crystallography 2010, 66 (Pt 4). 10.1107/S0907444910007493.

(68) Williams, C. J., Headd, J. J., Moriarty, N. W., Prisant, M. G., Videau, L. L., Deis, L. N., Verma, V., Keedy, D. A., Hintze, B. J., Chen, V. B., Jain, S., Lewis, S. M., Arendall, W. B., Snoeyink, J., Adams, P. D., Lovell, S. C., Richardson, J. S., Richardson, D. C. MolProbity: More and Better Reference Data for Improved All-Atom Structure Validation. Protein Sci 2018, 27 (1), 293–315. 10.1002/pro.3330.

(69) Berman, H., Henrick, K., Nakamura, H. Announcing the Worldwide Protein Data Bank. Nat Struct Mol Biol 2003, 10 (12), 980–980. 10.1038/nsb1203-980.

(70) Motulsky, H., Christopoulos, A. Fitting Models to Biological Data Using Linear and Nonlinear Regression: A Practical Guide to Curve Fitting; Oxford University Press, Incorporated: New York, UNITED STATES, 2004.

(71) Aurelio, L., Valant, C., Flynn, B. L., Sexton, P. M., Christopoulos, A., Scammells, P. J. Allosteric Modulators of the Adenosine A1 Receptor: Synthesis and Pharmacological Evaluation of 4-Substituted 2-Amino-3-Benzoylthiophenes. J. Med. Chem. 2009, 52 (14), 4543–4547. 10.1021/jm9002582.

(72) Miao, Y., Feher, V. A., McCammon, J. A. Gaussian Accelerated Molecular Dynamics: Unconstrained Enhanced Sampling and Free Energy Calculation. J. Chem. Theory Comput. 2015, 11 (8), 3584–3595. 10.1021/acs.jctc.5b00436.

(73) Jo, S., Kim, T., Iyer, V. G., Im, W. CHARMM-GUI: A Web-Based Graphical User Interface for CHARMM. Journal of Computational Chemistry 2008, 29 (11), 1859– 1865. 10.1002/jcc.20945.

(74) Tian, C., Kasavajhala, K., Belfon, K. A. A., Raguette, L., Huang, H., Migues, A. N., Bickel, J., Wang, Y., Pincay, J., Wu, Q., Simmerling, C. ff19SB: Amino-Acid-Specific Protein Backbone Parameters Trained against Quantum Mechanics Energy Surfaces in Solution. J Chem Theory Comput 2020, 16 (1), 528–552. 10.1021/acs.jctc.9b00591.

(75) Dickson, C. J., Walker, R. C., Gould, I. R. Lipid21: Complex Lipid Membrane Simulations with AMBER. J. Chem. Theory Comput. 2022, 18 (3), 1726–1736. 10.1021/acs.jctc.1c01217.

(76) Wang, J., Wolf, R. M., Caldwell, J. W., Kollman, P. A., Case, D. A. Development and Testing of a General Amber Force Field. J Comput Chem 2004, 25 (9), 1157–1174. 10.1002/jcc.20035.

(77) Case, D. A., Aktulga, H. M., Belfon, K., Cerutti, D. S., Cisneros, G. A., Cruzeiro, V. W. D., Forouzesh, N., Giese, T. J., Götz, A. W., Gohlke, H., Izadi, S., Kasavajhala, K., Kaymak, M. C., King, E., Kurtzman, T., Lee, T.-S., Li, P., Liu, J., Luchko, T., Luo, R., Manathunga, M., Machado, M. R., Nguyen, H. M., O’Hearn, K. A., Onufriev, A. V., Pan, F., Pantano, S., Qi, R., Rahnamoun, A., Risheh, A., Schott-Verdugo, S., Shajan, A., Swails, J., Wang, J., Wei, H., Wu, X., Wu, Y., Zhang, S., Zhao, S., Zhu, Q., Cheatham, T. E., Roe, D. R., Roitberg, A., Simmerling, C., York, D. M., Nagan, M. C., Merz, K. M. AmberTools. J Chem Inf Model 2023, 63 (20), 6183–6191. 10.1021/acs.jcim.3c01153.

(78) Roe, D. R., Cheatham, T. E. PTRAJ and CPPTRAJ: Software for Processing and Analysis of Molecular Dynamics Trajectory Data. J Chem Theory Comput 2013, 9 (7), 3084–3095. 10.1021/ct400341p.

