## Supplemental Data for "Structural basis of MK-97 positive allosteric modulation at the M_4_ mAChR"

**Table of Contents**

Supplementary Table1. 3

Supplementary Figures 1–5	4

Supplementary NMR, LC-MS, and HRMS data for MK-97	8

**Supplementary Table 1: Cryo-EM data collection, refinement, and model quality**

|  | M_4_ mAChR:ACh:MK-97:DNG⍺_si1_:Gβ_1_:Gɣ_2_:scFv16 |
| --- | --- |
| **Data Collection** |  |
| PDB code | 23OP |
| EMD code – Consensus | 69131 |
| EMD code – Focused  refinement of receptor | 69129 |
| Micrographs | 6129 |
| Electron Dose (e^-^/A^2^) | 53.6 |
| Voltage (kV) | 300 |
| Pixel size (Å) | 0.83 |
| Movie frames | 71 |
| Defocus range (µm) | 0.5 – 1.5 |
| **Refinement** |  |
| Symmetry imposed | C1 |
| Particles (final map) | 347,823 |
| Resolution @0.143 FSC (Å)*  Consensus  Receptor local refinement | 2.7  2.55 |
| CC_map–model_ (volume) | 0.86 |
| **Model Quality** |  |
| R.M.S. deviations |  |
| Bond length (Å) | 0.004 |
| Bond angles (^o^) | 0.722 |
| Ramachandran |  |
| Favoured (%) | 98.2 |
| Outliers (%) | 0 |
| Rotamer outliers (%) | 0 |
| C-beta deviations (%) | 0 |
| Clashscore | 3.53 |
| MolProbity score | 1.14 |

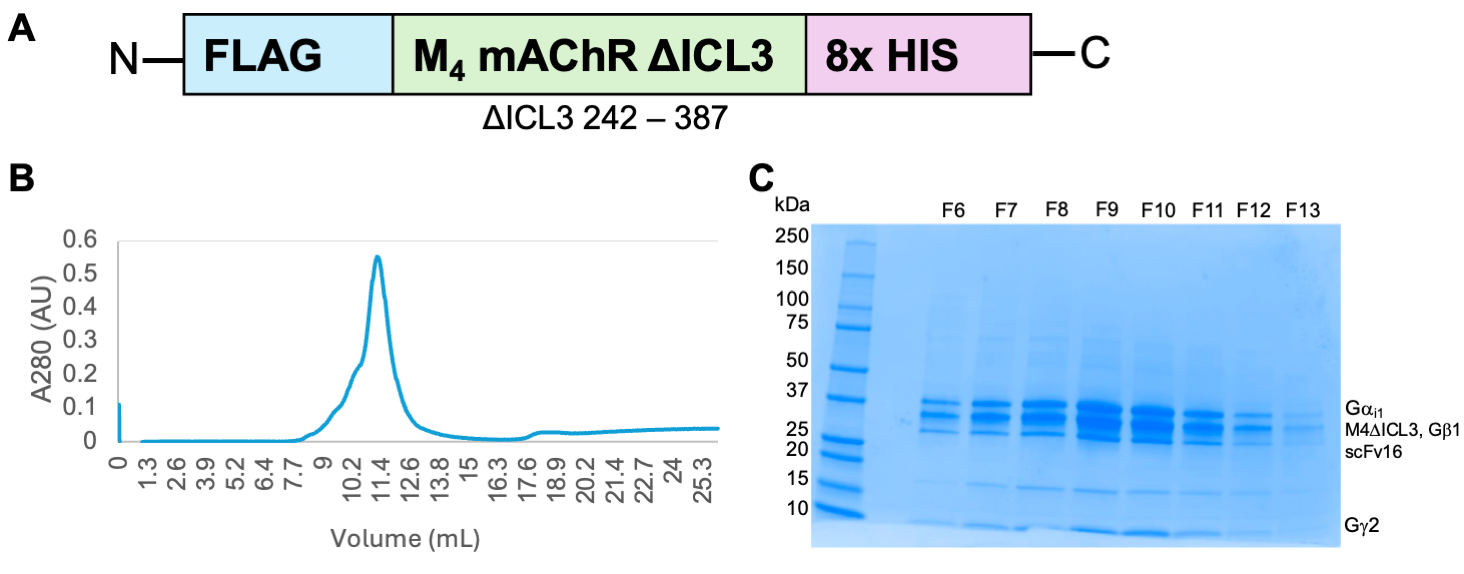
**Supplementary Figure 1: Purification of M_4_R:ACh:MK-97 complex**

(**A**) Construct map of the M_4_ mAChR used in this study. An N-terminal FLAG epitope tagged human M_4_∆ICL3 (residues 242 – 387 deleted) with a C-terminal 8X-Histidine tag and a 3C protease cleavage site. (**B**) Size exclusion chromatography trace of the purified M_4_R:ACh:MK-97:DNG⍺_i1_:Gβ_1_Gɣ_2_:scFv16 complex. (**C**) Coomassie stained SDS-PAGE of the components from the purification.

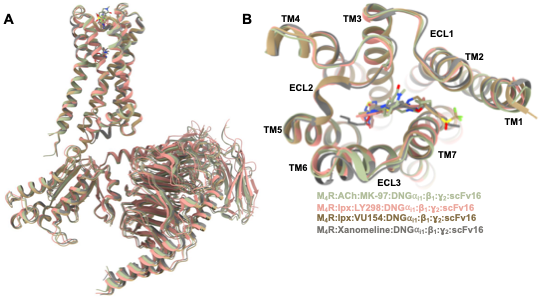

**Supplementary Figure 2: Structural comparison of agonist-bound M_4_ mAChRs determined by cryo-EM.**

(**A**) Overall view of the M_4_ mAChRs complexed to DNGα_i1_ bound to ACh and MK-97, shown in green, iperoxo, and LY2033298 (PDB: 7TRP) shown in pink, iperoxo and VU0467154 (PDB: 7TRQ) shown in brown, and xanomeline (PDB: 8FX5) shown in grey. (**B**) Extracellular view comparing ECLs and TM regions.

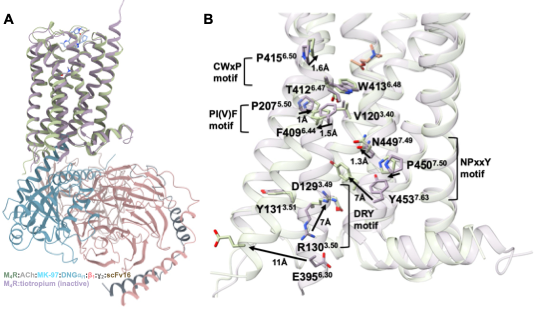

**Supplementary Figure 3: Comparison of the inactive and ACh, MK-97-bound active states of the M_4_ mAChR complex**

(**A**) Comparison of the tiotropium-bound M_4_ mAChR model (PDB:5DSG) and the active M_4_ mAChR complex model, shown as cartoons purple and green, respectively. (**B**) Membrane view of residues, shown as sticks, and conserved motifs involved in activation of class A GPCRs.

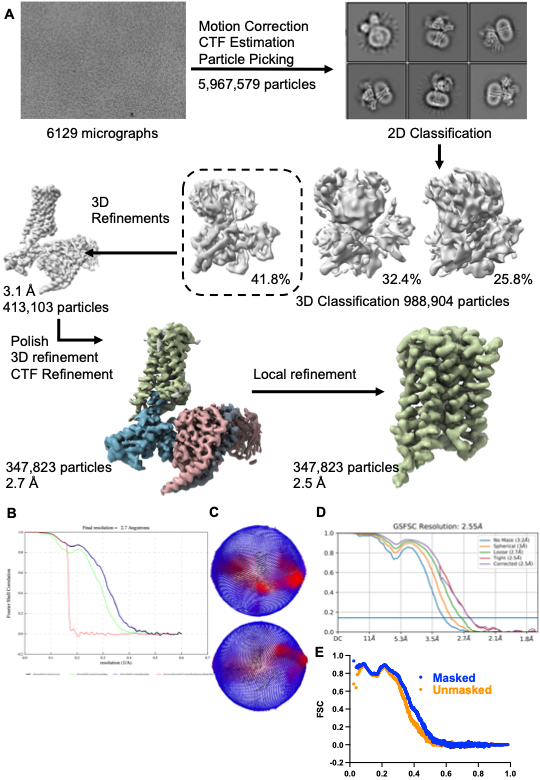

**Supplementary Figure 4: Cryo-EM processing for the MK-97-bound M_4_ mAChR G protein complex**

**(A)** Cryo-EM data processing workflow as described in methods. Gold-standard Fourier shell correlation (GSFSC) plot for the consensus map **(B)** and **(D)** the receptor focused map. **(C)** The angular distribution of particles projected in the final map. (**E**) FCS map-to-model fit.

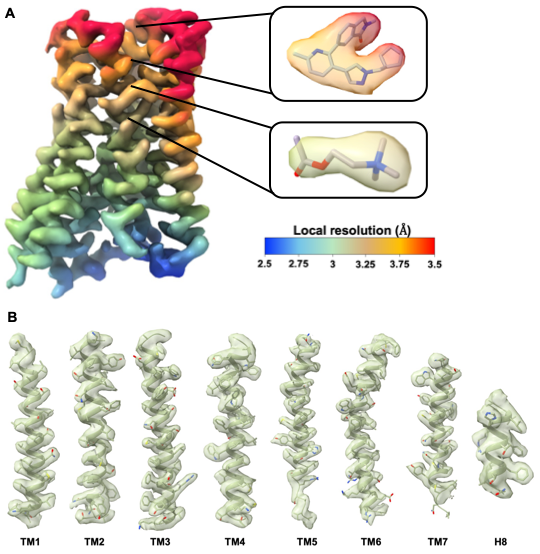

**Supplementary Figure 5: Cryo-EM maps**

(**A**) Local resolution of the receptor-focused refinement map with corresponding local resolution of ACh in the orthosteric site and MK-97 in the allosteric site of the M_4_ mAChR. (**B**) Cryo-EM map (contour level 0.22) from the consensus map and modelling for the 7 transmembrane helices and helix 8 of the receptor.

**NMR, LC-MS and HRMS data confirmed >95% purity for MK-97:**

**6-(3-(1-(Cyclopentylmethyl)-1*H*-pyrazol-4-yl)-6-methylpyridin-2-yl)-2-methylisoindolin-1-one**

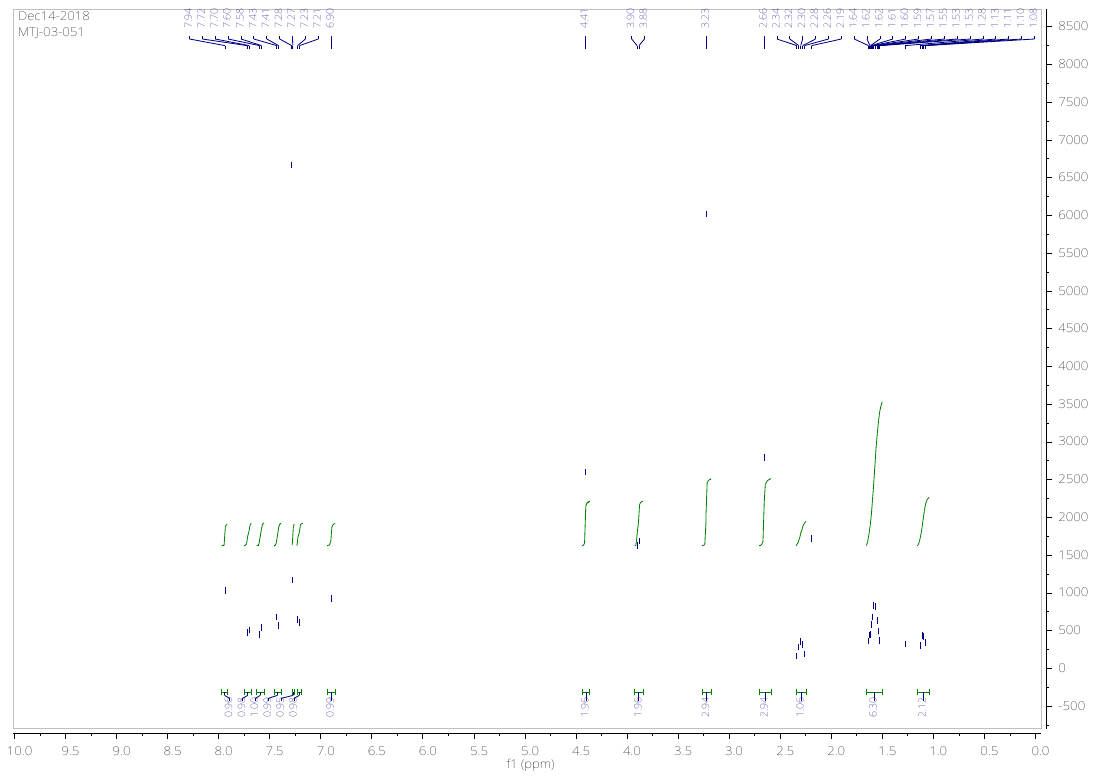

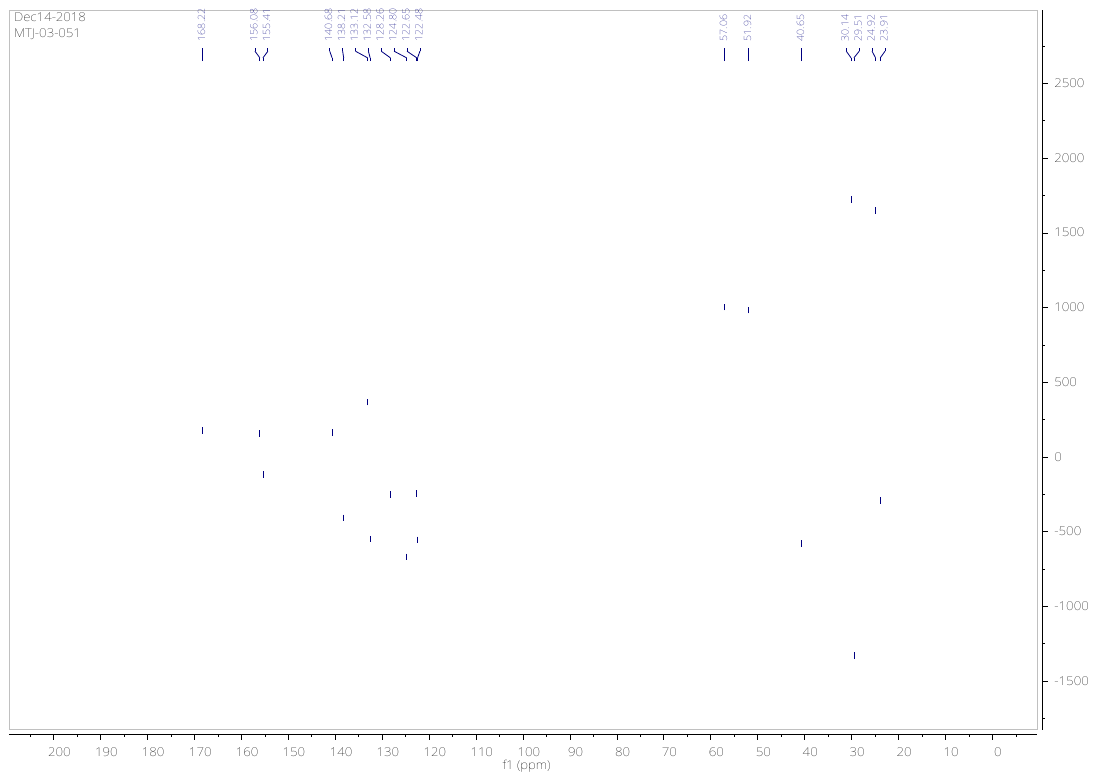

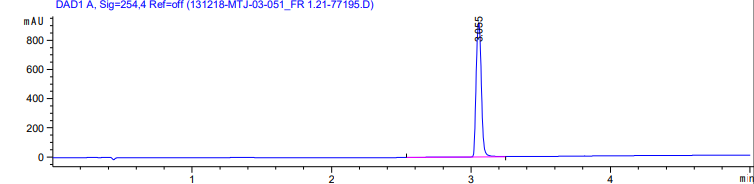

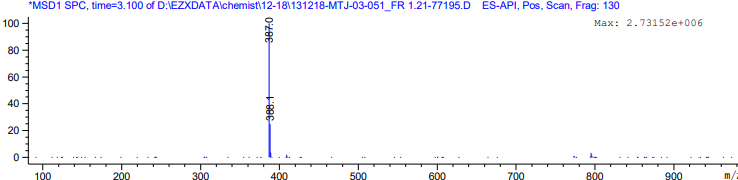

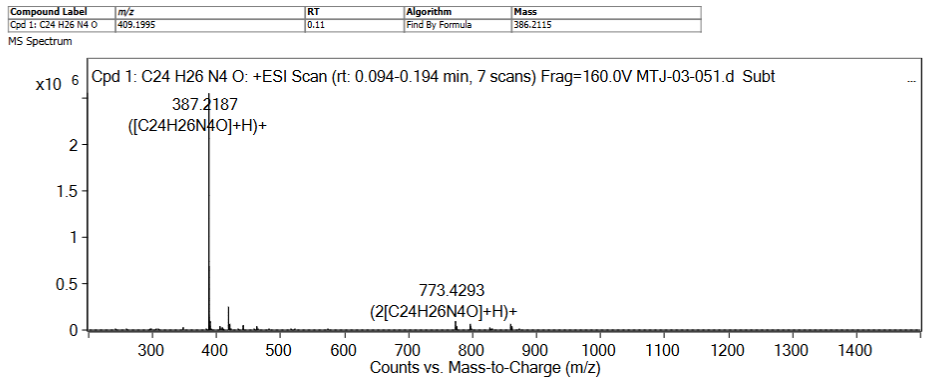

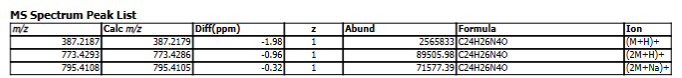
